# Deleterious Mechanical Deformation Selects Mechanoresilient Cancer Cells with Enhanced Proliferation and Chemoresistance

**DOI:** 10.1101/2022.07.22.501206

**Authors:** Kuan Jiang, Su Bin Lim, Jingwei Xiao, Doorgesh Sharma Jokhun, Menglin Shang, Xiao Song, Pan Zhang, Lanfeng Liang, Boon Chuan Low, G.V. Shivashankar, Chwee Teck Lim

## Abstract

Cancer cells derived from secondary tumors can form new distant metastases more efficiently as compared to their primary tumor counterparts. This is partially due to the unfavorable microenvironments encountered by metastasizing cancer cells that result in the survival of a more metastatic phenotype from the original population. However, it is unclear how cancer cells may acquire such metastatic competency after overcoming deleterious mechanical stresses. Here, by forcing cancer cells to flow through small capillary-sized constrictions, we demonstrate that mechanical deformation can select a tumor cell subpopulation that exhibits resilience to mechanical squeezing-induced cell death. Transcriptomic profiling reveals up-regulated proliferation and DNA damage response pathways in this subpopulation, which are further translated into a more proliferative and chemotherapy-resistant phenotype. Our results highlight a potential link between the microenvironmental physical barriers and the enhanced malignancy of metastasizing cancer cells which may potentially be utilized for novel therapeutic strategies in preventing the metastatic spread of cancer cells.

## 1. Introduction

Metastasis is a rare event despite the constant detection of a few circulating tumor cells (CTCs) in the peripheral blood of cancer patients^[1,2]^. Fluid shear stress^[3,4]^, immune surveillance^[5]^, and the small-sized capillaries^[6,7]^ can effectively destroy the majority of metastasizing tumor cells entering the bloodstream. Secondary tumors are found to have different biophysical^[8]^ and molecular characteristics^[9]^ compared to their primary counterparts, and they are often more lethal and impose great challenges to cancer treatment. Thus, elucidating the mechanism of metastasis and finding potential methods to stop the metastatic spread is an important research topic.

Recent studies have shown that the physical microenvironment of tumor cells plays an important role in the progression of cancer^[10,11]^. For example, mechanical stresses arising from tight tissue space may rupture the nuclei of metastasizing cancer cells that squeeze inside and increase the genome instability of cancer cells^[12–14]^. Meanwhile, the mechanical stimuli from the shear force in the bloodstream can promote cancer cell survival in circulation, and enhance extravasation and their resistance to chemotherapy drugs^[15,16]^. This evidence suggests that the physical forces encountered during the metastatic journey of cancer cells may significantly modulate the metastasizing cancer cells. Additionally, the metastatic cascade can be considered an evolutionary process that selects specific traits of cancer cells that are capable of overcoming different deleterious stresses^[17–19]^. For example, the tight squeezing and mechanical stresses arising from geometry confinement in tight tissue space or capillaries have been shown to damage and eliminate most of the metastasizing tumor cells^[6]^. It is widely known that metastatic cancer cells are generally more invasive and exhibit resistance to cancer therapies compared to the localized primary tumor cells, however, it is unclear whether the mechanical stresses play a direct role in this transformation. More specifically, it is important to answer whether there exists a specific cancer subpopulation that is resilient to mechanical stresses and what characteristics the remaining population exhibits after mechanical stress cleared those non-resilient cells.

To tackle this problem, we explored the possibility of selecting a cancer cell subpopulation that is resilient to mechanical stress in this study. Comparing such a selected subpopulation and the non-selected counterpart provides a novel cellular model to explore the factors influencing cell survival under mechanical stresses and how the population after such selection differs from the original population. Such a model is free from any artificial genomic or proteomic perturbations, and thus can reflect the molecular signatures at their endogenous level with only the influence of a transient mechanical challenge. The findings associated with the resilient subpopulation provide novel connections on how the mechanical stresses may shape the metastasizing cancer cells and the workflow and experimental model can be readily adopted for more broadly exploring the metastasis under physical influence.

## 2. Results

### 2.1 Resilience to deformation is a selectable trait

We first designed a workflow to pick out the cancer cells that can survive after deforming through constrictions with dimensions similar to that of the capillaries^[20]^. To perform a high throughput deformation, we developed and optimized a microfluidic deformation assay that can perform such a mechanical selection on 30,000 suspended cancer cells per minute (Figure. S1a-g, movie S1, see Materials and Methods section). To determine whether the survivor cancer cells after the harsh mechanical deformation are inherently resilient to such deformation, we performed a two-step deformation experiment using the microfluidic deformation assay (Figure. 1a) for H1650 (lung cancer cell line), MCF7 (breast cancer cell line) and MDA-MB-231-Luc (luciferase-expressing MDA-MB-231 metastatic breast cancer cell line, abbreviated as MDA-Luc) cells. The results showed that most of the cells that survived the first deformation (the survivor cells) also survived the second deformation (Figure. 1b) suggesting the existence of a cancer subpopulation that is resilient to deformation which we term “mechanoresilient cells” here. We subsequently collected and expanded the survivor cells after passing the original cancer cell population (ORI) through the microfluidic deformation assay for one round (Figure. 1c, this process is termed “mechanical selection” in this study). With different cancer cell lines, we found the mechanoresilient phenotype is expandable in MCF7, Mia-PaCa2, MDA-Luc, and BT549-Luc (luciferase-expressing BT549 breast cancer cell) cell lines (Figure. 1d). Although the survivor H1650 cells showed resilience to deformation in the two-round deformation experiment, such phenotype disappeared after cell expansion while the expanded survivor CaCo2 cells show no resilience to deformation as well. As MCF7 is an epithelial non-metastatic breast cancer cell line from the very upstream of the metastasis cascade, it is of particular interest to explore whether and how the mechanoresilient subpopulation exists in these early-stage cancer cells. Thus, we chose MCF7 as the subject for a more in-depth investigation. We first performed multiple rounds of the mechanical selection on the cells where we found one round of selection (the selected cells are subsequently termed S1 cells) is enough to maximize such a selection effect (Figure. 1e, 22.87%±7.31% in ORI vs. 52.30%±8.12% in S1 MCF7, mean±s.d.). It is noteworthy that we, at the same time, found such resilience to deformation-induced cell death would gradually disappear with time (Figure. S1h) in the population. However, the time window of the mechanoresilient phenotype was long enough for subsequent cellular and molecular characterizations. These results combinedly showed the possibility of selecting and expanding a novel cancer subtype featuring resilience to mechanical deformation-induced cell death. These selected cancer cells together with the unselected original population can form an excellent cell model to study the factors that are crucial for cell survival under extreme mechanical stresses.

**Figure. 1.**
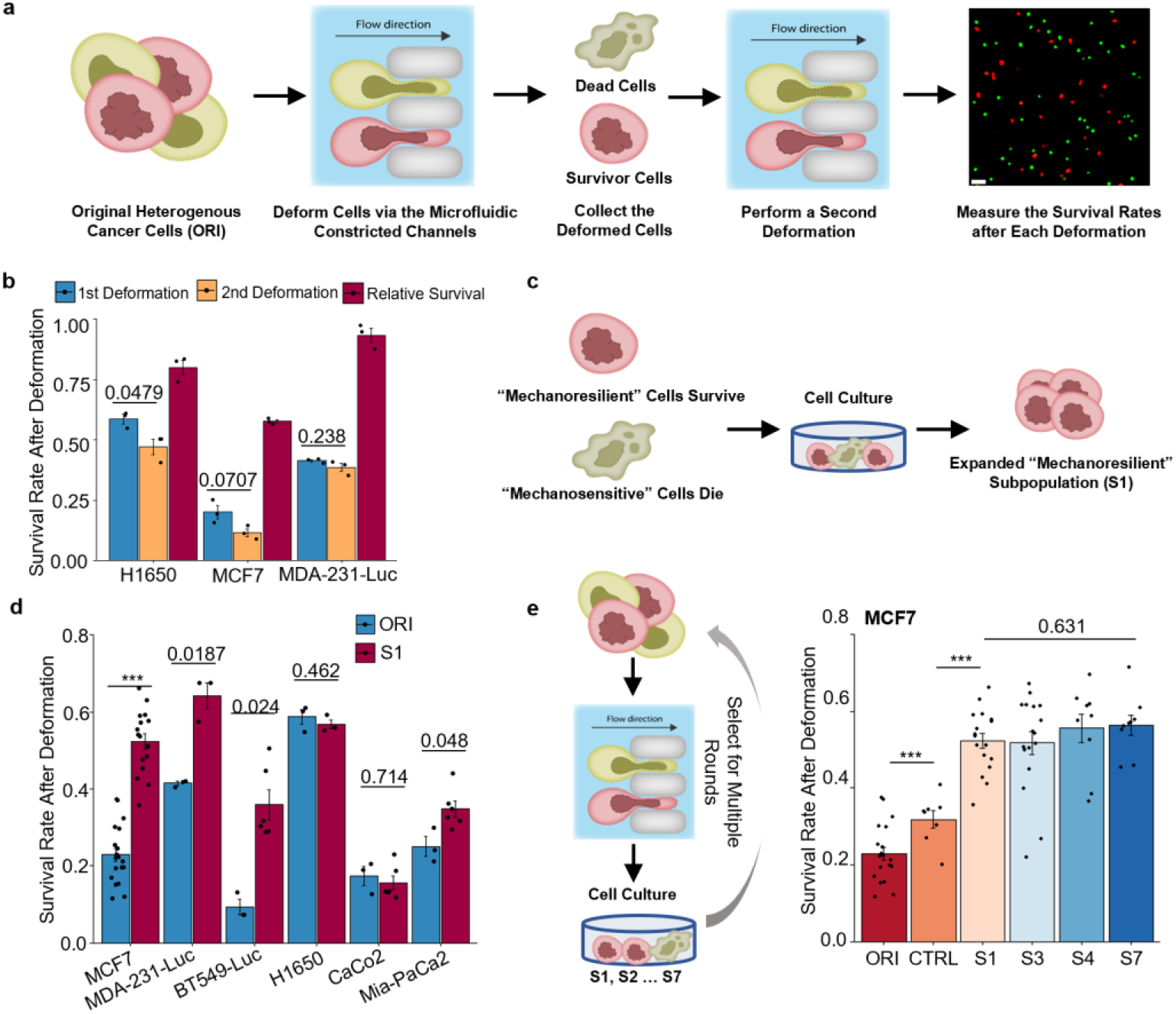
Mechanical deformation selects mechanoresilient cancer cells. **(a)** Schematics of using a microfluidic deformation assay to perform two rounds of mechanical deformation and measure the survival rate after each round of deformation. **(b)** The survival rate of H1650, MCF7, and MDA-MB-231-Luc cells after each round of deformation. The relative survival is calculated by dividing the 2^nd^ round survival by the 1^st^ round survival. Barplot shows mean± s. e.m.; unpaired, two-tailed, t-test, exact p-value labeled for p>0.01; **: p<0.01;***p<0.001. n=3, each jitter shows one independent experiment, the criteria are consistent throughout the manuscript, details can be found in the method section. **(c)** Schematics of expanding the survivor cells after deforming the ORI cells through the microfluidic deformation assay. **(d)** The survival rate of the expanded survivor cells after deforming through the microfluidic deformation assay compared to the respective ORI cells. Barplot shows mean± s. e.m.; MDA-Luc and H1650: unpaired, two-tailed, t-test; the rest: Wilcoxon rank-sum test; each jitter shows one independent measurement. **(e)** Left: Schematic of a multiple-round mechanical selection experiment performed on MCF7 breast cancer cells. Right: Quantification of the survival rate of the ORI MCF7 cells and selected MCF7 cells (S1, S2… indicating rounds of selection, CTRL are the cells selected with 20 μm wide constrictions) after passing through the microfluidic deformation assay, each jitter showing one independent measurement; statistics on ORI vs.CTRL and CTRL vs. S1: Wilcoxon rank-sum test; on multiple comparisons on S1, S3, S4, S7: one-way ANOVA; each jitter shows one independent experiment.

### 2.2 Cellular characteristics associated with the mechanoresilience

By comparing the size distribution of MCF7 selected for different rounds to the original population, we found that cell size is not a determining factor in the survival after deformation (Figure. S2a). To further confirm this, we took advantage of the size difference when MCF7 cells are seeded in different cell densities, where we found the differently sized MCF7 cells indeed show no survival difference (Figure. S2b). As cell deformability is considered an important factor influencing how cancer cells cope with mechanical stresses, we then used a microfluidic micropipette assay to compare the deformability of ORI and S1 MCF7 cells. However, we did not observe any significant difference in the cellular level deformability in this study as well (Figure. S2c-e, movie S2).

While performing the selection experiment on different cell lines, we found there are two distinct types of cellular responses to the deformation. For CaCo2 and Mia-PaCa2 cells, there were significant cellular ruptures when the cells passed through the constrictions, which were not seen in H1650, MCF7, MDA-MB-231, and BT549 cells (Figure. S3a and c). This resulted in a significant cell loss during the deformation process for Mia-PaCa2 and CaCo2 (from 2.34×10 ^5^ in the original cell suspension to 0.51×10 ^5^ in the collected cell suspension leading to a 78.2% cell loss), while the cell loss for the non-rupturing cell lines was only about 3%. The cell loss in the 2^nd^ round deformation of Mia-PaCa2 cells was about 54.9% which was similar to the expanded S1 population, indicating a selection effect on the cellular ability to resist cell ruptures in this specific cell line (Figure. S3b). By examining videos of the deformation of a non-rupturing cancer cell line MCF7, only a small portion of cancer cells (<1%) experienced cellular rupture during the deformation^[21]^ (Figure. S3d-e, movie S3). This indicated that the ability to maintain cellular integrity during deformation is an important factor influencing survival but was not the major contributor to the observed mechanoresilience in the MCF7 cells.

It has been extensively reported before that an intact cell nucleus is essential in determining cell survival after extreme nuclear deformation^[12,14,22,23]^. Indeed, we found that a large portion of the dead cells after deformation exhibited ruptured nuclei with DNA extruded out from the nuclear lamina (Figure. 2a-c). Ruptured nuclei were rare in the survivor cells, however, we observed locally accumulated lamin A/C or nuclear blebbing in the survivor cells 30 minutes after deformation (Figure. S4a-c), which is consistent with the previously reported nuclear lamina rupture and repair after constricted migration^[24]^. This also indicates that nuclear rupture can nevertheless happen in the survivor cells but to an extent that is repairable ^[13,22]^. In the previously reported confined migration scenario, nuclear deformation was relatively slow, and cells could have enough time to repair a ruptured nucleus, but here the flow-driven deformation process was much faster and the cells with significant nuclear rupture were unlikely to be able to repair themselves. Thus, limiting the nuclear lamina rupture during the deformation should be a key factor for the cancer cells to survive in our setting. To test this, we first investigated the expression of lamin A/C and lamin B1, which are two essential structural proteins supporting the nuclear lamina^[22,25]^, in MCF7 cells before and after the mechanical selection. Compared to the ORI population, S1 MCF7 cells had a significantly higher level of lamin B1 and a slightly higher level of lamin A/C (Figure. 3a). To study the correlation between lamin expressions and cell survival after deformation, we analyzed the ORI and S1 cells stained with a live/dead marker-propidium iodide (PI)-immediately after deformation. This allowed us to distinguish the survivor cells from dead cells right after the selection. From the immunostaining, we found the survivor cells typically exhibited a significantly higher level of lamin B1 while the level of lamin A/C did not show a consistent correlation with survival (Figure. 2d-f). The lamin A/C level here also excludes the possibility that the observed low lamin B1 in dead cells is a result of protein degradation after cell death. This high lamin B1 phenotype was also found in the survivor cells of MCF7-Luc, MDA-Luc, BT549-Luc, H1650, and Mia-PaCa2 after deformation (Figure. 4a-f). Additionally, in the MDA-Luc cells, the subpopulation with a lamin B1 intensity in the lower 25^th^ percentile had only an 11.3% survival rate, in contrast to the 62.4% survival rate in the upper 75^th^ percentile (Figure. 4g). Meanwhile, in all tested cell lines, the lamin B1 intensity in most of the survivor cells is above the 70^th^ percentile (Figure. 4h). Knockdown of lamin B1 in both ORI and S1 MCF7 cells reduced the survival rate after deformation significantly (Figure. 3b-c). Note that we did not see apparent nuclear blebbing in LMNB1-KD MCF7 cells as well which has been previously reported in some cell lines after lamin B1 knockdown (Figure. S4d)^[26,27]^. We subsequently found lamin B1 overexpression could elevate the survival rate of MDA-Luc and BT549-Luc cells after deformation (Figure. 4i). This evidence suggests that lamin B1 plays a critical role in determining cell survival after deformation. To gain a deeper insight into how lamin B1 and lamin A/C might behave differently during deformation, we transfected MCF7 cells with mCerulean-lamin B1 and mCherry-lamin A/C. Surprisingly, we found lamin A/C could be transiently segregated from a compressed nucleus, and the intensity of lamin A/C along the nuclear periphery was decreasing during the deformation. Lamin B1 was intact during the entire deformation process before being forced to rupture in a 2μm constriction (Figure. S4e-f, movies S4, S5). These distinct dynamics of lamin A/C and lamin B1 suggested that lamin B1 primarily protected the nuclear lamina integrity during a rapid nuclear deformation in MCF7 breast cancer cells.

**Figure. 2.**
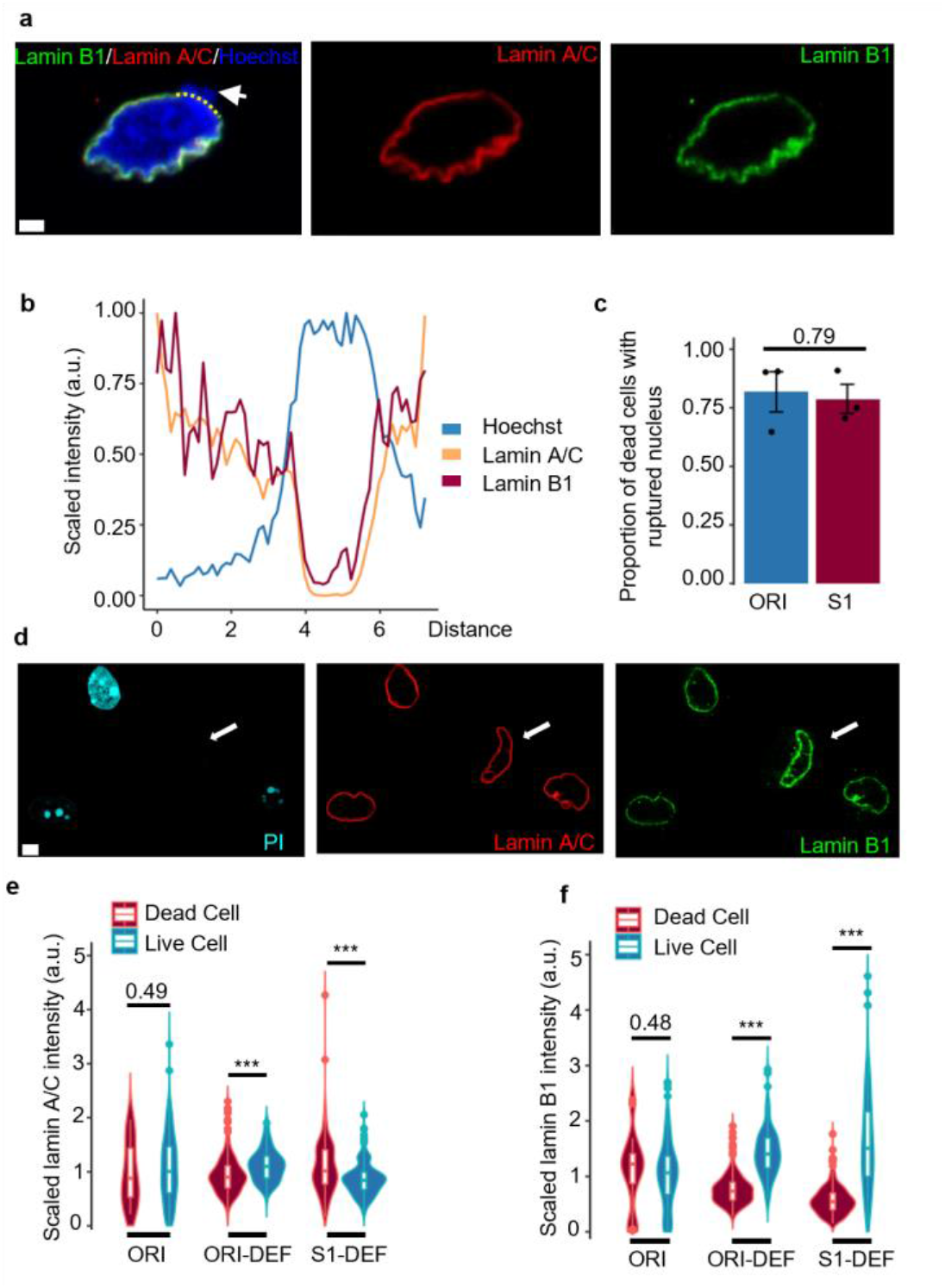
Lamin B1 protects mechanoresilient MCF7 from deformation-induced nuclear ruptures and cell death. **(a)** Representative images of the nucleus of a dead MCF7 cell after deformation, the white arrow pointing to the site of lamina rupture with DNA leaking out from the nucleus; the yellow dashed line marked the line for the intensity profile plotted in **(b)**; Scale bar: 2μm. **(b)** The intensity profile of DNA (Hoechst stained), lamin A/C, and lamin B1 along the line of the nuclear lamina, the intensity values are scaled to (0,1]. **(c)** Quantification of the percentage of ruptured nuclei in the dead cells after deformation; t-test; Data from 3 independent repeats of each group, total of 110 cells in ORI and 37 cells in S1. **(d)** Representative images of the relative lamin abundance with Live/Dead labeling (PI stained) of cells after deformation: live cells (PI negative) show significantly higher lamin b1 expression as indicated by the white arrows; Scale bar: 5μm. (**e-f**) Quantification of the relative (scaled to the mean of each group) lamin intensity in dead cells and live cells after deformation, ORI denotes original MCF7 cells that are not deformed through the microchannels with naturally apoptosis cells; Wilcoxon rank-sum test; results are from 3 independent repeats in each group, total >200 cells were imaged and quantified in each group.

**Figure 3.**
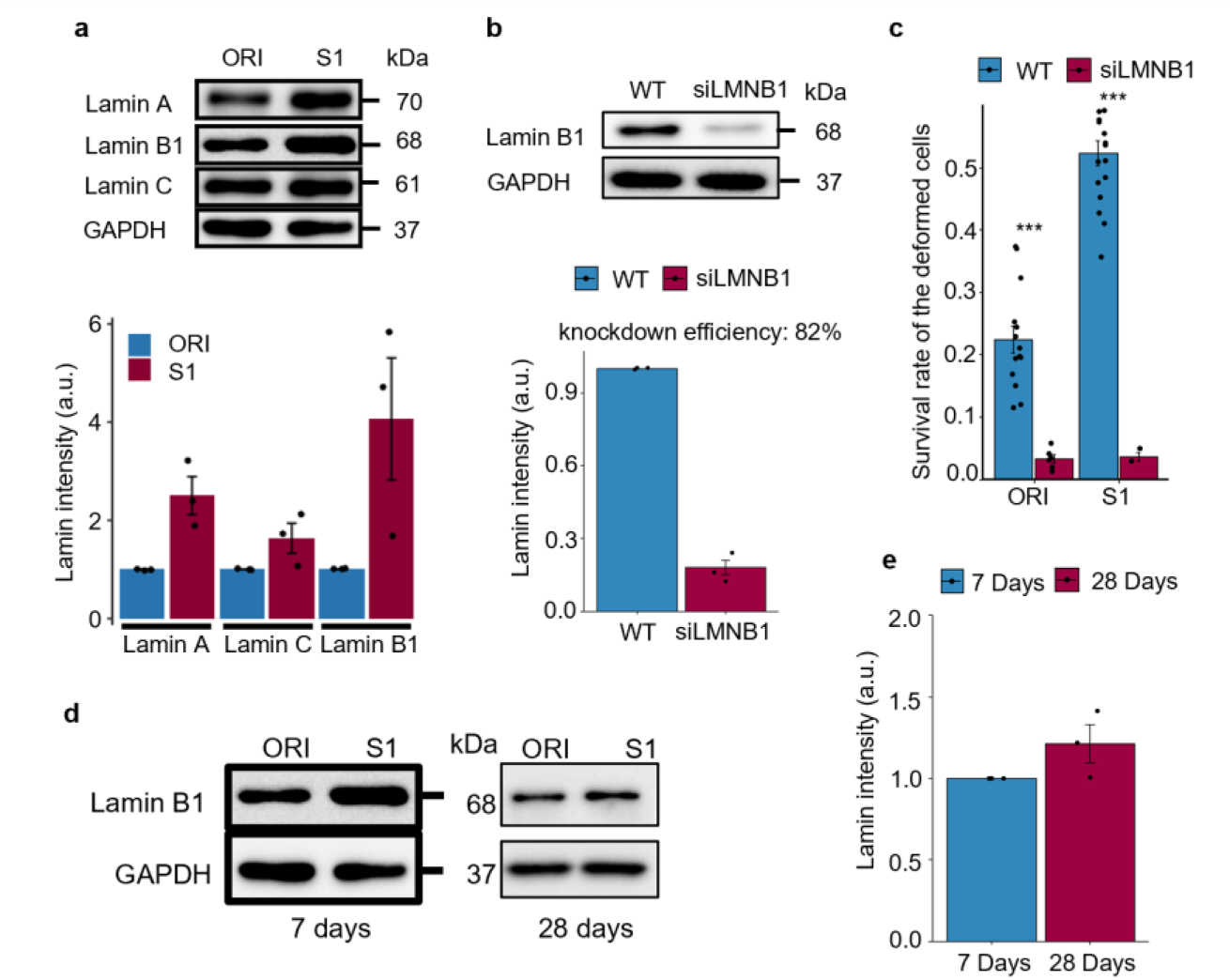
Quantification of lamin protein levels and their influence on survival after deformation. **(a)** Western blot of lamin A, lamin C, and lamin B1 in ORI and S1 cells, with the quantification (below) comparing the relative intensity of the bands; Quantification showing results from 3 independent repeats of each group. S1 cells are lysed at the first passage (7 days) when reaching confluency. The relative expression is calculated by dividing the S1 lamin level by its corresponding ORI group. **(b)** Western blot of lamin B1 after transfecting MCF7 cells with LMNB1 RNAi for 48 hours; the quantification of knockdown efficiency is performed on ORI cells. **(c)** Quantification of the survival rate of LMNB1 knockdown MCF7 cells after deformation, 3 independent knockdowns were performed in each group, n=547 cells in ORI-KD and n=591 cells in S1-KD were quantified. **(d)** Comparison of the lamin B1 level of S1 cells 7 days and 28 days after selection; the 7 days result is from panel **(a). (e)** Quantification of the relative lamin B1 level on day 28 compared to day 7; n=3.

**Figure. 4.**
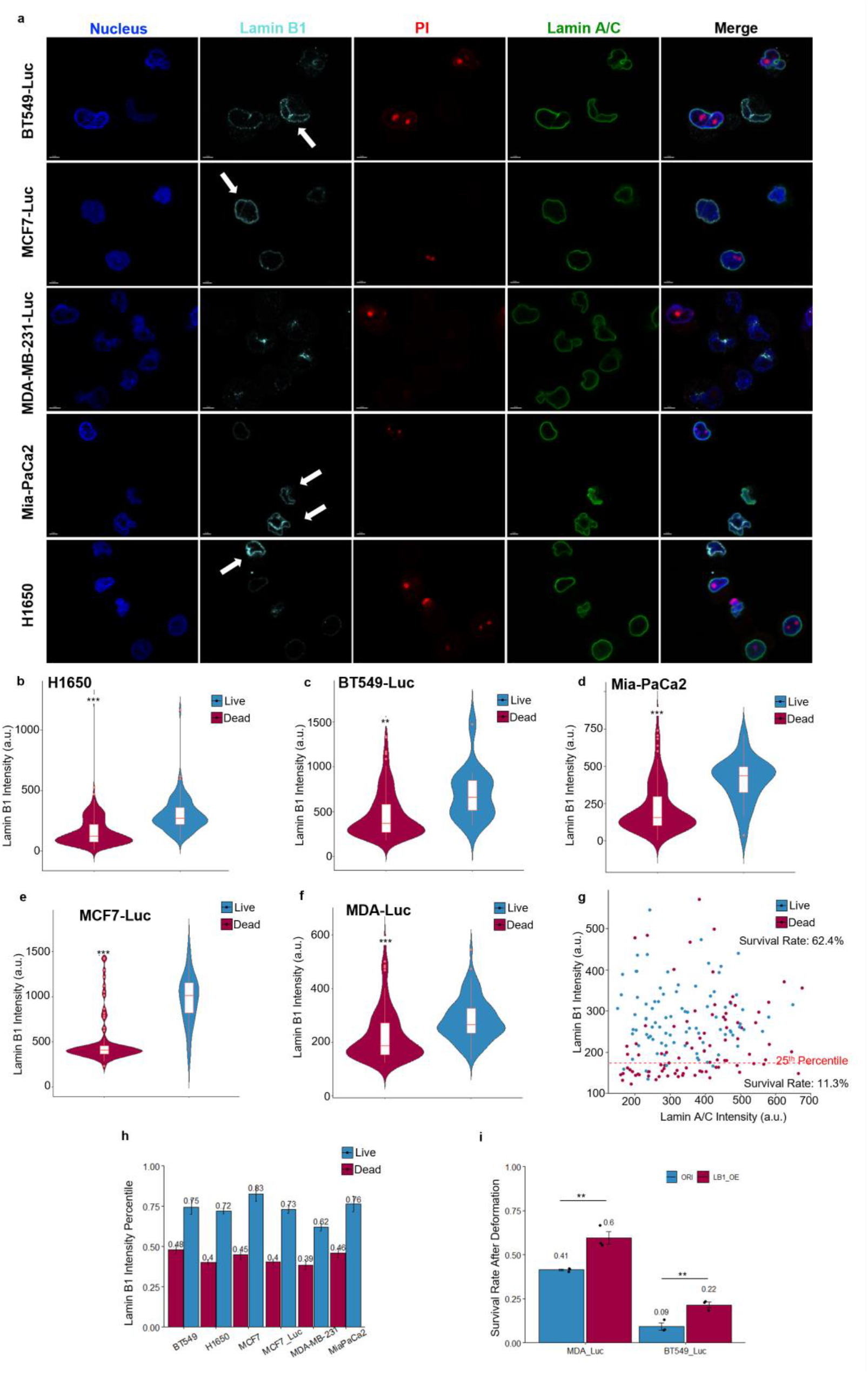
Survivor cells after mechanical selection have high lamin B1 levels. **(a)** Representative immunofluorescence images of the survivor cells in MCF7-Luc, H1650, Mia-PaCa2, MDA-Luc, and BT549-Luc stained for HOECHST (nucleus), PI (live/dead); lamin A/C, and lamin B1. Arrows showing the surviving high lamin B1 cells in the respective cell lines after the mechanical deformation. **(b-f)** Quantification of lamin B1 intensity in live and dead cells of each tested cell line; Wilcox rank-sum, 3 independent samples for each cell line, at least 100 cells are quantified in each group. **(g)** Co-plot of lamin A/C and lamin B1 intensity in deformed MDA-Luc cells. The dotted red line shows the 25th percentile cutoff of lamin B1 intensity. The survival rate for cells below this cutoff is 11.3% and 62.4% above the cutoff. **(h)** Quantification of the lamin B1 percentile in the live and dead population of each cell line after deformation. Barplot showing mean±s. e.m. **(i)** The survival rate of lamin B1 overexpressed MDA-Luc and BT549-Luc after deformation; n=3; unpaired, two-tailed, t-test.

To understand whether there were other changes in the nuclei of selected cells, we characterized the nuclear structures of ORI and S1 MCF7 cells at different cellular states. The nuclear sizes of the survivor cells were found slightly smaller than that of the dead cells (Figure. 5a-e). We subsequently found that the nuclear lamina was originally wavy and invaginated^[28,29]^ in the suspended state (Figure. 5b,f) but was fully stretched out during deformation (Figure. 5g). This is consistent with a recent observation of the nuclear lamina unfolding under compression which was reported as an essential mechanosensing process^[30]^. Apart from this, we also observed that the S1 MCF7 cells compared to the ORI cells, while having a similar projected nuclear area and perimeter in the suspended state, had a larger nuclear lamina surface area (Figure. 5h), and a larger projected area and perimeter in the adherent state (Figure. 5i-k). As the nuclear lamina is known to be stretched by the cytoskeletal forces during cell spreading^[31]^ (Figure. 5i), these results indicated that there was a more reserved nuclear lamina surface area in the S1 cells, which might buffer the strain during nuclear stretching as the cells passed through the constrictions. Taken together, this evidence suggested that higher lamin B1 expression and larger nuclear lamina surface areas might limit cell death during extreme cell deformation which contributed to a mechanoresilient breast cancer subpopulation after mechanical selection.

**Figure. 5.**
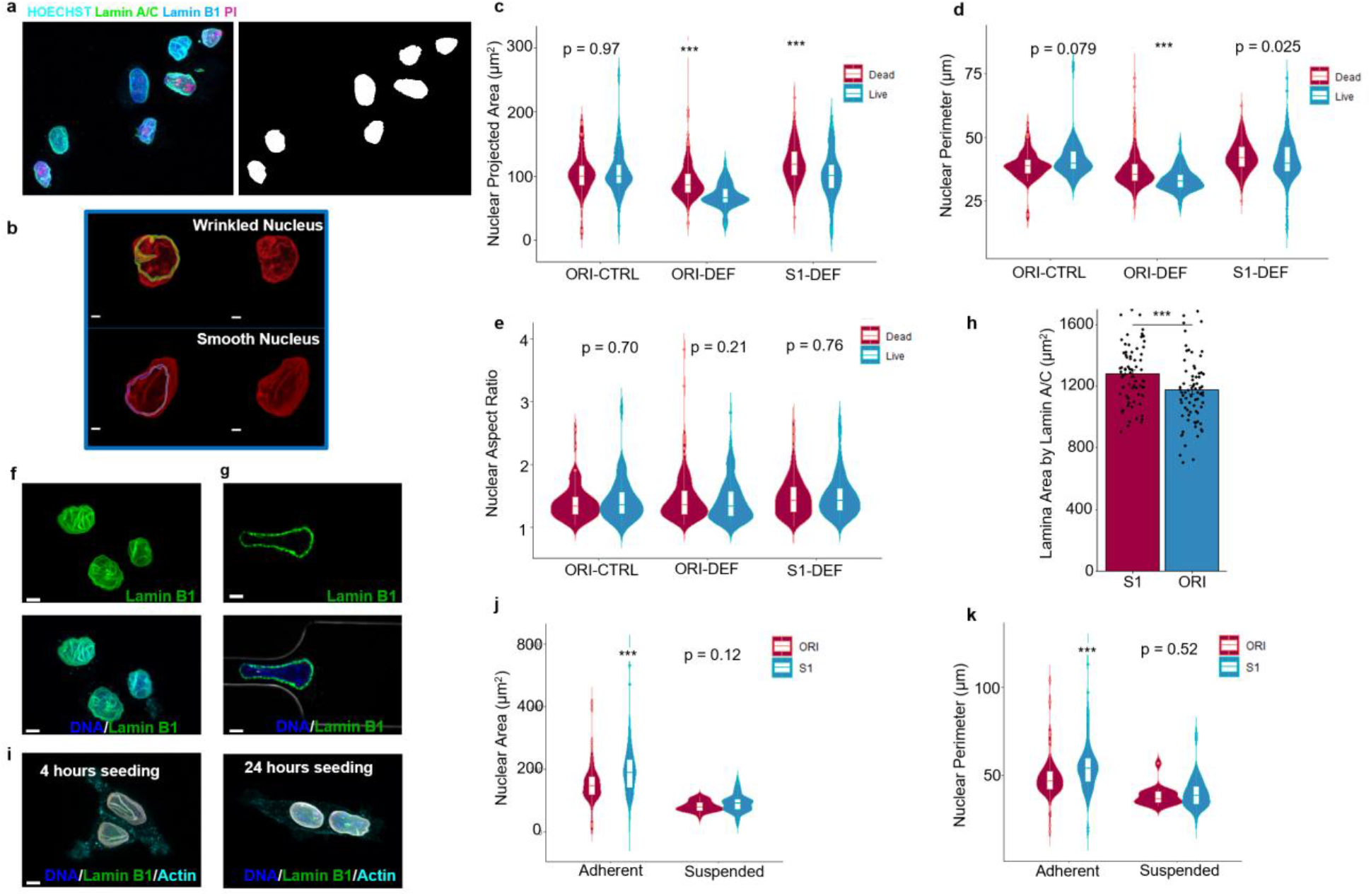
Comparing the nuclear phenotypes of the original and mechanoresilient breast cancer cells. **(a)** A representative image of a nucleus after deformation stained with PI for labeling the live/dead cells. **(b)** Reconstructed nuclear envelope from lamin A/C immunostaining. A wrinkled nucleus shows deep invagination of its nuclear lamina. Scale bar: 2μm. **(c)** Comparison of the projected nuclear areas, **(d)** perimeters, and **(e)** aspect ratios in the MCF7 cells with or without selection with live/dead labeling, at least 100 nuclei are analyzed from 3 independent repeats in each group; Wilcoxon rank-sum tests. **(f)** Suspended nuclei, and **(g)** a stretched nucleus during deformation stained with lamin B1. The wavy nuclear lamina gets stretched out during deformation. Scale bar: 4 μm. **(h)** Comparison of the nuclear surface areas determined by lamin A/C, Wilcoxon rank-sum test, n=3, jitters showing data distribution. **(i)** Lamin B1 labeled nuclear lamina 4 hours or 24 hours after seeding, the wavy invaginated lamina gets stretched out after the cells fully spread. **(j)** comparing the areas and **(k)** the perimeters of the nucleus at adherent or suspension states in the ORI or S1 MCF7 group. Wilcoxon rank-sum tests; at least 100 cells from 3 repeats are quantified.

To gain an insight into how lamin gene expressions are associated with cancer development, we did a pan-cancer meta-analysis in the TCGA database^[32]^. First, we compared the expression of LMNA, LMNB1, and LMNB2 in normal and primary tumor tissues (Figure. S5a-c). Among the 24 cancer types investigated, we found both B-type lamins: lamin B1 and lamin B2 showed significant enrichment in tumor tissues (22/24 and 23/24 tumor types investigated, respectively), but LMNA was only enriched in 9/24 cancer types (Figure. S5d-g). There was a slight downregulation of LMNA in breast tumor tissues while LMNB1 and LMNB2 were significantly upregulated. These results indicated a strong correlation between B-type lamin and cancer development. Prognostic analyses further revealed that LMNA, LMNB1, and LMNB2 expression all had prognostic value in the distant metastasis-free survival (DMFS) of breast cancer patients (Figure. S5h-j).

### 2.3 Enhanced proliferation and DNA damage resistance after mechanical selection

We next sought to obtain a more comprehensive understanding of the transcriptomic changes in the mechanically selected cancer cells. ORI, S1, and a flow control group (CTRL, see Materials and Methods section) MCF7 cells were sent for RNA sequencing (Figure. S6a). Only 6 downregulated and 19 upregulated genes were found differentially expressed (adjusted P < 0.05 and |Log2FoldChange| >0.5) between ORI and CTRL cells (Figure. S6b), indicating a transient exposure to high shear stress is not causing significant transcriptomic changes in the MCF7 cells. A total of 479 and 280 differentially expressed genes (DEGs) were identified when comparing the S1 vs. ORI or S1 vs. CTRL groups, confirming that the mechanical selection resulted in a significantly altered gene expression profile in the S1 cells (Figure. S6b, c). To focus on the effect of mechanical deformation, we performed functional enrichment analyses on the S1 vs. CTRL DEG list. Cell proliferation and DNA damage response (DDR) pathways were found to be the top influenced cellular functions after mechanical selection (Figure. S6d-e). A hallmark gene set enrichment analysis (GSEA) of the DEGs (Figure. S6f) revealed the targeted genes of two proto-oncogenes, E2F (enrichment score [ES] = 0.66, p < 0.001) and MYC (ES = 0.59, p < 0.001), are significantly upregulated in the selected MCF7 cells, which further implied enhanced tumorigenesis ability of the mechanoresilient cancer cells^[33,34]^.

We subsequently performed two proliferation assays to investigate whether the altered proliferation genes were translated to a more proliferative phenotype. Indeed, although the selected cancer cells showed an initial cell cycle delay similar to previous reports^[35,36]^(Figure. 6a, b), we found the expanded (>7days post-selection) S1 MCF7 cells exhibited a much higher proliferation rate compared to the ORI cells (Figure. 6c, d). We went on to test whether the altered DDR gene expression had any functional implications. First, to investigate whether altered DDR in S1 MCF7 cells implicated a DNA damage resistance, we treated the S1 MCF7 cells with a chemotherapy drug doxorubicin (Dox) which is known to cause DNA double-strand breaks (DSBs)^[37]^. While the half-maximal inhibitory concentration (IC50) value was not significantly changed, the selected cells exhibited much higher viability as characterized by MTT assays at a high Dox concentration (Figure. 6e). To test whether the cells exhibit resistance to DNA damage induced by Dox, we evaluated the DSB level in ORI and S1 MCF7 cells after being treated with 1μM Dox which was shown to induce high cell mortality in Figure. 6e. The S1 cells treated with 1μM Dox for 12 hours had a reduced γH2Ax intensity as well as lower γH2Ax foci numbers, indicating a chemotherapy resistance at high drug concentration (Figure. 6f-h).

**Figure. 6.**
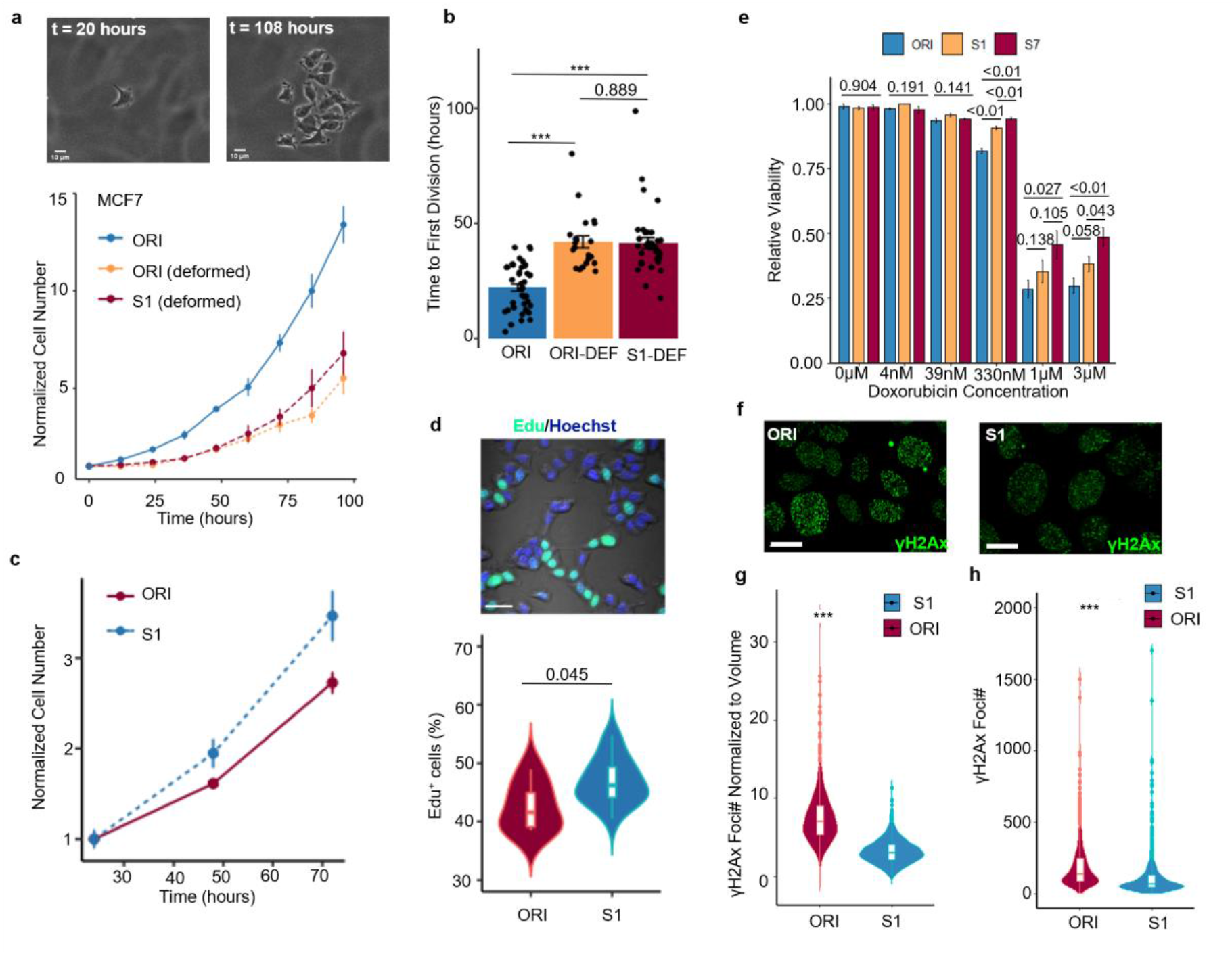
Enhanced proliferation and DNA damage repair ability in mechanoresilient MCF7 cells. **(a)** Top: Representative cell tracking images for quantifying the cell proliferation right after selection. Scale bar = 10 μm. Bottom: Quantification of the cell proliferation right after mechanical selection, the point range is showing the normalized mean number (to the first time point) ± s. e.m. at different time points after cell seeding. **(b)** Quantification of the time to the first cell division after cell seeding comparing the ORI MCF7, the deformed ORI, and the deformed S1 MCF7 cells; three experiments, ORI:41 cells, ORI-DEF: 22 cells, S1-DEF: 36 cells are quantified; p values are calculated with Wilcoxon rank-sum test. **(c)** Quantification of the tracked cell number at different time points after seeding (normalized to the seeding density of the respective groups), point range showing scaled cell number ± s. e.m. **(d)** Top: A representative image of the Edu stained nuclei, and, Bottom: quantification of the Edu positive cell percentage comparing the ORI and S1 MCF7 cells, Wilcoxon rank-sum test, ORI: 6 repeats; S1: 15 repeats. **(e)** Quantification of cell viability after being treated with different concentrations of Doxorubicin, point range showing mean±s. e.m.; for 330nM, 1μM, and 3μM: unpaired, two-tailed, t-test; for the multiple comparisons: one-way ANOVA; n=3 in each condition. **(f)** Representative images of γH2Ax staining in ORI (left) and S1 (right) MCF7 cells treated with 1μM Doxorubicin. **(g)** Quantification of the γH2Ax numbers per unit volume and **(h)** the γH2Ax Foci number per nucleus; Wilcox rank-sum test, 3 independent groups in each condition.

### 2.4 DEGs after mechanical selection implied a poor disease prognosis

The DEGs in the mechanically selected cells provided a panel of genes that may potentially bridge the intravascular mechanical selection to the malignant metastatic phenotype. Moving forward, we used the DEGs that were common in S1 compared to both ORI and CTRL populations for a meta-analysis in a large, combined breast cancer cohort (Figure. S6g-h). Approximately 40% of the identified DEGs demonstrated predictive power in at least one of the clinical endpoints analyzed (relapse-free survival (RFS), overall survival (OS), and distant metastasis-free survival (DMFS); Figure. 7a-c). The prognostic genes were most prominently associated with proliferation, suggesting that the altered proliferation pathways after mechanical selection are related to poor breast cancer prognostics (Figure. 7d). By analyzing the paired primary tumor and metastasis tumor RNA-Seq dataset in the TCGA BRCA cohort, we found 7 common genes that express differently in the metastasis tumors as well as in S1 MCF7 cells (Figure. 7e-k), and 31 common genes after removing the log2foldchange cutoff in the DEG list (Figure. S7a). Additionally, we found a score given by a weighted expression of these 31 genes (see Materials and Methods section) is a strong prognostic marker of breast cancer (Figure. S7b-d), which further implied that the breast cancer cells have changed toward a more malignant subtype after the mechanical selection.

**Figure. 7.**
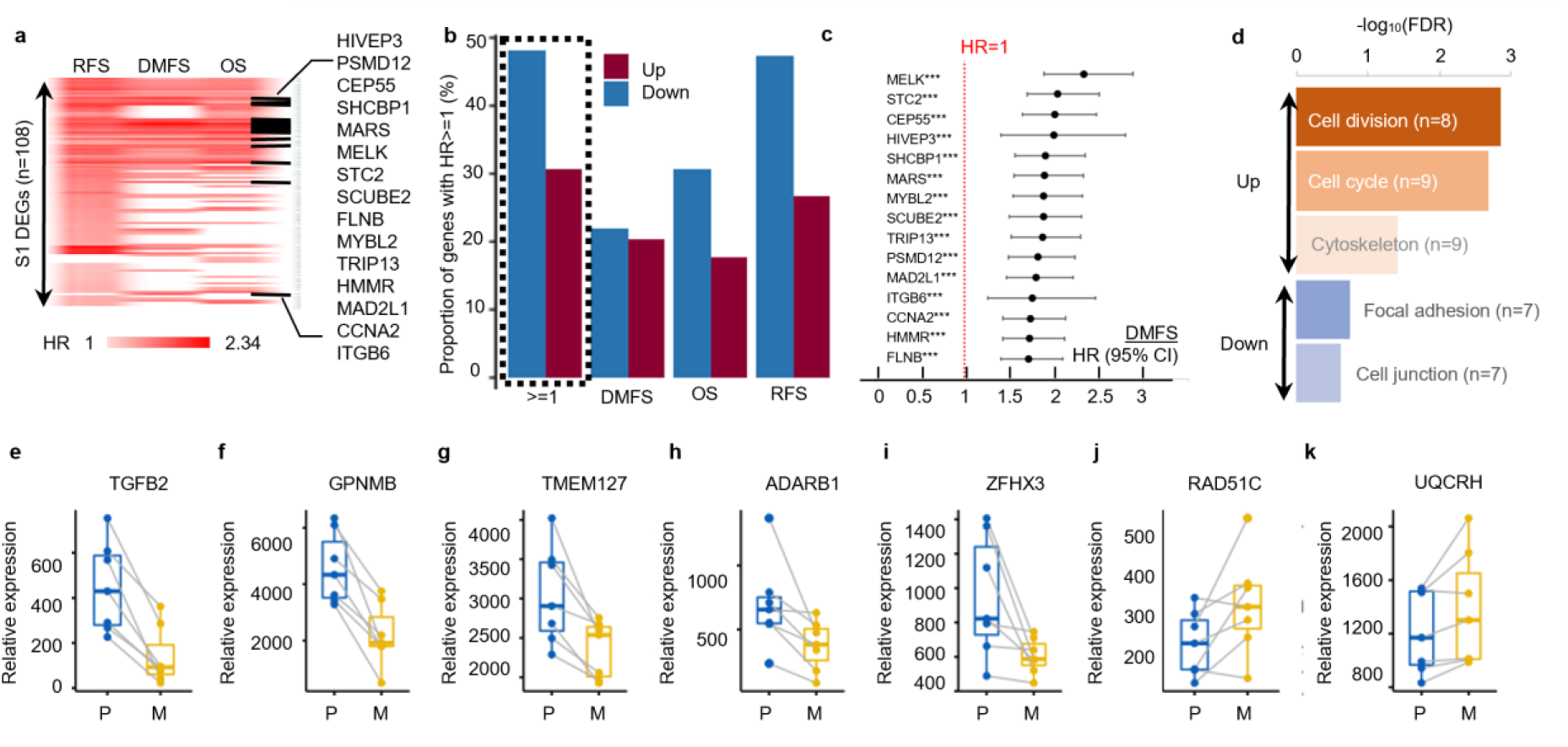
Mechanoresilience gene signatures are correlated with breast cancer metastasis and poor clinical prognosis. **(a)** Heatmap showing the common DEGs in ORI vs. S1 and CTRL vs. S1 DEG lists that have prognostic power in at least one of the clinical endpoints: RFS (relapse-free survival), DMFS (distant metastasis-free survival), and OS (overall survival); the color bar shows the hazard ratio derived from survival analysis; for downregulated genes with HR<1 the values are reverted; black dash line labels genes having prognostic power in all three endpoints. (**b)** Quantification of the percentage of total up/down-regulated genes that have predictive power in each clinical endpoint. (**c)** Forest plot of top genes with HR>1.7 in the DMFS of breast cancer patients. (**d)** Top enriched functional clusters in the list of genes with prognostic value, the colors in functional cluster names indicate down or upregulated function and the color bar shows the levels of FDR. (**e-k)** Expression levels of the DEGs overlapping in the CTRL vs. S1 group and the paired primary tumors and metastasis tumors from TCGA BRCA cohort, P: primary tumor; M: metastasis tumor.

Finally, we analyzed the mechanoresilient gene signatures in an earlier work that performed *in vivo* selection on MDA-MB-231 metastatic breast cancer cells for lung metastasis (MDA_LM2 in GSE2603)^[18]^. We found a total of 61 genes that are overlapping between the MDA_LM2 DEGs and MCF7_S1 DEGs (Figure. S8a-b). The weighted expression score of these 61 genes demonstrated strong prognostic power in the 3-year lung metastasis-free survival (LMFS) time and marginal prognostic power in the 5-year LMFS of the breast cancer patients in the same study cohort (Figure. S8c, d). The score also seemed to have a strong correlation with the survival of patients with lymph node metastasis at diagnosis (Figure. S8e). This 61-gene panel was also found strongly prognostic in a larger breast cancer cohort with 1,764 patients (Figure. S8f-h). Altogether, these 61 genes formed a putative panel that represented the mechanical aspect of an *in vivo* metastasis evolution and was correlated with poor breast cancer prognosis.

## 3. Discussion

The physical barrier to metastasis is an important innate defense of the human body against tumor invasion, and the understanding of how such barriers become ineffective during cancer progression is a key to making malignant tumors a localized disease. Although mechanical stresses are thought to destroy most of the metastasizing cancer cells^[38]^, the survivor cells are of major interest in understanding the key factors influencing cell survival under extreme mechanical stresses. Here, we established a mechanical selection workflow using microfluidic chips which can be conveniently used for selecting cancer cells that may survive after a deleterious mechanical deformation. Through this study, we proved the existence of a novel cancer cell subtype - mechanoresilient cancer cells – featuring the resistance to mechanical deformation-induced cell death. This generated mechanoresilient subpopulation is free from any laboratory gene or protein perturbations and can be specifically used to study the influence of mechanical stress on cancer cells. The cell model formed through such a mechanical selection process can be used to further investigate what kind of a cancer cell is likely to survive under deleterious mechanical stress. It should be noted here that one limitation of using this methodology is the possible coupled effect of mechanical selection and mechanical modulation. The altered molecules between the original population and the selected subpopulation may be due to the survival advantage of the subpopulation or the mechanical stress altered the relevant molecular pathways in the survivor cells. The live/dead staining method used in this study can mostly resolve this issue as the short time scale of the selection process is generally inadequate for the activation of protein expression as the cells are fixed immediately after deformation. Simultaneously staining structurally similar proteins (e.g. lamin A/C and lamin B1 in this work) can further exclude the influence of protein degradation due to cell death. This method, however, may be insufficient to exclude the possibility of a further upregulation/downregulation of the protein of interest in the expanded cell population. Such a potential combined effect can be decoupled through single-cell tracking after the selection.

By using this model, we examined the association between cellular and nuclear size and survival after deformation. The nuclear size in the survivor population is smaller compared to the dead population, which is intuitive as smaller nuclei are expected to experience less damage during the deformation. This is not contradicting the similar cell size observed in the MCF7 population selected for different rounds or the lack of survival difference of MCF7 cells cultured to different sizes. One reasonable explanation is that the survival advantage of MCF7 cells with smaller cell sizes or smaller nucleiwas insignificant compared to the more dominant factors revealed in this study.. A recent study also found the aspect ratio of cells after deformation might indicate the metastatic potential of cells^[44]^. In our case, we had a similar observation that the more metastatic MDA-MB-231 cells exited the channel with a higher and more prolonged aspect ratio. We confirmed from the MCF7 cells that the survivor cells were not showing different nuclear aspect ratios after the selection, but whether the cellular aspect ratio had any implication in survival needs further investigation. The nuclear lamina was found with a larger area in the selected cells after spreading. Given the observation that the nuclei had a higher reserved area in the suspended state, we expect their larger spread area was dominantly contributed by such reserved area before spreading. However, it should be noted that this does not exclude the possibility that a higher cytoskeleton contraction force contributes to the larger lamina area after spreading.

Our result suggested a high lamin B1 expression is strongly associated with cell survival after deformation, which is consistent with a recent study showing that lamin B1 could limit nuclear rupture in migrating neurons lacking lamin A/C^[39]^. One arising question is how lamin B1 mechanistically associates with the observed resilience to deformation. First of all, the survivor cells after deformation should inherently have a higher expression level according to our results.

The observed association is unlikely due to a mechanically induced lamin B1 upregulation considering that the cancer cells are fixed immediately after the deformation. Thus, the high lamin B1 level in the expanded S1 population of MCF7 cells, in our opinion, is also more likely due to the originally high lamin B1 level in the survivor cells. It should be noted that the lamin B1 level in the selected cells would gradually decrease during extended cell culture. As discussed earlier, at this point, we were not able to excludethat the high lamin B1 phenotype in the expanded S1 MCF7 cells might be additionally influenced by a short-term mechanical activation of lamin B1 expression. However, another possibility is that the high lamin B1 phenotype is unstable and can only exist transiently in the MCF7 population from the beginning, which resulted in the dynamic lamin B1 level we observed here^[40]^. Although lamin B1 was shown to be strongly associated with cell survival after deformation of the different cell lines, it should also be noted that the tested cell lines have vastly different lamin B1 baseline expressions. For example, BT549 and MDA-MB-231 used in this study have a lower lamin B1 expression compared to the MCF7 cells. However, MDA-MB-231 exhibited a higher survival after deformation compared to MCF7 while BT549 showed the opposite. An accompanying observation is that the tested cell lines had shown distinct phenotypes of their cell body integrity after deformation (see Figure S3). Meanwhile, all cell populations consistently showed that their higher lamin B1 expressed cells exhibited a significantly higher survival rate after deformation. Altogether, these results suggest that lamin B1 is one common factor that influences the resilience to deformation in all tested cell lines.

Another observation is the different deformation dynamics of lamin A/C and lamin B1 during deformation. It seems that lamin A/C is more mobile in the nuclear lamina and can be segregated from the nucleus under large nuclear strain in MCF7 cells which potentially may be due to its different structural organization^[45,46]^ of the lamin B1 in the lamina. Although our findings emphasize mainly on lamin B1 rather than lamin A/C, this does not contradict previous findings on the roles of lamin A/C in confined migration^[22]^ or the loss of lamin B1 in promoting lung cancer metastasis^[41]^. Our findings highlighted here are that the function of different lamin subtypes in different stages of cancer progression is context-dependent. While lamin A/C has long been found to regulate nuclear mechanics^[42,43]^, these results together with our findings on the association between lamin B1 and the observed mechanoresilience added new insights into a previously undermined role of B-type lamins in regulating the integrity of nuclear lamina during extreme deformation and the progression of cancer^[32]^.

Through RNA sequencing and functional characterization, we also observed an unexpected connection between the mechanical selection process and the enhanced proliferation as well as the activated DNA damage response. Although the difference in MTT determined viability of ORI and S1 MCF7 cells seems marginal in the doxorubicin concentration test, the S7 MCF7 cells which have been challenged by the mechanical stress for multiple rounds show significant higher viability at high drug concentration. Additionally, despite the insignificant viability difference compared to ORI cells, S1 MCF7 shows a great reduction in DNA damage level after being treated with high concentration of doxorubicin which was shown to cause a high reduction in cell viability. This is a compliment to the recent findings in the cancer cell migration under confinement^[12]^. Circulating tumor cells and the cancer cells in secondary tumors are found to be more tumorigenic and resistant to DNA damaging chemotherapy drugs. Our finding provided a clue that the coping to mechanical stress in the metastatic cascade may contribute to the gain of proliferation ability and drug resistance. Unlike the previously reported mechanically-induced proliferation through mechanosensitive channels^[47]^, the altered proliferation and DNA damage response observed here is a result of a transient deformation which have not been reported before. Cellular lamin B1 expression was previously found to associate with telomere and chromosome instability^[48,49]^, recruitment of DNA damage repair proteins^[50]^, and cellular senescence^[51]^. However, similar to the resilience to deformation, the elevated lamin B1 level reverted with time in our study while the enhanced proliferation could remain in the selected cancer cells for more than 2 months (Figure. 3d-e), which suggested the proliferation change here may be independent of the high lamin B1 phenotype. Furthermore, the transcriptomic level changes brought by the selection process are found to indicate a poorer prognostic in breast cancer patients. In a summary, the mechanical stress in the metastatic cascade is selecting and altering the metastasizing cancer cells and plays a crucial role in transforming cancer cells to the hard-to-treat metastatic phenotype.

Metastasis is a long process that drives the primary tumor cells to evolve towards a more lethal form^[52,53]^. We have shown here a new aspect that a deleterious mechanical deformation can potentially be a driving force for such a metastatic evolution. The detailed mechanisms underlying the selection process and how the lamina structure influences cell survival after deformation is yet to be further studied and we expect the relevant findings can be translated to inhibit metastatic spread with our innate physical microenvironment. We believe future studies building on our findings to further reveal the mechanisms underlying such a selection process can lead to novel therapeutic options that limit cancer to a treatable local disease.

## 4. Materials and Methods

### Cell culture

All cell lines used in this study were purchased from ATCC, grown, and maintained at 37ºC with 5% CO _2,_ and routinely tested for mycoplasma contamination. Breast cancer cell lines *MCF7* and *MDA-MB-231* were cultured in DMEM (Lonza) supplemented with 10% FBS (Gibco) and 5μg/ml gentamycin (Gibco). Both cell lines were passaged weekly with medium refreshing every two days. All cells were grown to 90% confluency before subculturing or used for experiments. The culture medium used for *MCF10A* was MEGM (Lonza, with SingleQuots™ Kit), for the *H1650* lung cancer cell line, *MDA-MB-231-Luc, BT549-Luc* (kind gift from George Yip’s group) was RPMI 1640 (Lonza) supplemented with 10% FBS and 5μg/ml gentamycin, for *CaCO2* was MEM with 20% FBS, 100IU/ml penicillin and 100mg/ml streptomycin (Pen-Strep), and for Mia-Paca2 was DMEM with10% FBS, 1% nonessential amino acids, 1% sodium pyruvate and 1% glutamine. For each independent repeat of mechanical selection, the ORI cells were cultured in different flasks in parallel to account for the batch effect. Before experiments, cells were detached from cell culture flasks by removing the culture medium, rinse with DPBS 3 times, and treated with 0.05% trypsin-EDTA (0.1x of the original concentration) for 10 minutes in a 37 ºC incubator. After the incubation, enzyme digestion was stopped by adding an equal volume of the corresponding complete culture medium, and cells were spun down with a centrifuge at 200g for 3 minutes. The cell pellets were collected and resuspended in the cell culture medium or experiment buffer for subsequent usages. All cell lines are authenticated by the manufacturer. MCF7 and MDA-MB-231 were additionally authenticated with RNA sequencing results. All cell lines did not belong to the list of commonly misidentified cell lines.

### Transient transfection

mCherry-Lamin A/C (Addgene #55068) and mCerulean-Lamin B1 (Addgene #55380, both plasmids were kind gifts from Tony Kanchanawong’s lab) were used for studying the deformation dynamics of lamin A/C and lamin B1 in MCF7. Cells were seeded to 80% confluence in T25 flasks and transfected with jetPrime (Cat#114-15) for 24 hours before harvesting them for experiments. mCerulean-Lamin B1 was also used for the lamin B1 overexpression experiment with 2.5μg DNA/well in a 6-well plate. A note here is we tried several different lamin B1 constructs and found overexpression of lamin B1 can cause significant cell death in MCF7, probably due to the mitotic catastrophe in high lamin B1 cells. RNAi for LMNB1 knockdown (MISSION esiRNA, Merck Cat#EHU057911) or scramble control RNAi is transfected (jetPrime) for 48 hours in 80% confluent MCF7 cells seeded in T25 flasks per manufacturer’s instruction. Knockdown efficiency was confirmed through the western blot of lamin B1 protein.

### Microfluidic devices

Microfluidic devices were fabricated using PDMS soft lithography^[54]^ with SU8 photoresist features on a silicon wafer. In brief, the design of each chip was drawn with AutoCad 2019 (Autodesk) and printed on a sodalime photomask. The designed features were then printed to SU8 by UV exposure (all master wafers were fabricated by the microfabrication core in Mechanobiology Institute). After receiving the master wafers, the wafer surfaces were briefly treated with oxygen plasma at 15W, 8.8sccm for 30s (Tergeo, PIE Scientific), and silanized (1H, 1H, 2H, 2H-perfluorooctyl trichlorosilane) for 2 hours in vacuum. The silanized mold was then used for device fabrication with Polydimethylsiloxane (PDMS, Sylgard 184 Elastomer Kit, Dow Corning) at a ratio of 10:1. The PDMS was poured onto the mold after mixing well and degassed for 1 hour until all bubbles had disappeared. The device was subsequently baked at 70 ºC for at least 2 hours. The resulting PDMS devices were cut with a razor blade and peeled off carefully, punched with 1.5mm holes at inlets and outlets, and bonded to glass slides after both surfaces were activated with oxygen plasma for 30s. The final devices were left in a 70 ºC oven overnight for strengthening t he bonding before using them for experiments.

### Microfluidic setup optimization

There were two microfluidic assays used in this study: the microfluidic deformation assay and the microfluidic flow control assay (fig. S1A). The only difference between the two assays was the gap between pillars (5μm in the deformation assay and 20μm in the control assay). The survival rate of MCF7 cells passing through different sized constrictions is shown in fig. S1C. Very few cancer cells survived after deforming through a 3μm×10μm (cross-section) constriction. The survival rate using one long or multiple short constrictions was similar (fig. S1D). For better mimicking the periodic deformation that circulating tumor cells might experience in different *in vivo* settings which were considered the most deleterious scenario *in vivo*^[6,55,56]^ and considering the technical robustness of microfabrication, we adopted the design with multiple short constrictions. We initially tested the infusion mode with both constant pressure (pneumatic pump) and constant volume flow rate (syringe pump). The stress-induced on cancer cells (reflected by survival rate, fig. S1E-G) is similar in both scenarios, however, the constant volume flow rate reached a much higher throughput (not sensitive to channel clogging). Thus, we used a constant volume flow rate for our microfluidic system.

### Generation of mechanoresilient cancer cells

Microfluidic devices with dimensions of 10μm × 5μm × 20 (heigh×width×rows) and with 126 constricti ons in each row were used for generating the mechanoresilient cancer cells in this study (schematically shown in fig. S1A). The devices fabricated using the aforementioned methods were flushed with 70% alcohol for 1 min at 100μl/min before flushing by DPBS with 0.01% Pluronic F-68 (Sigma) for 1 min at 100ul/min. The chip was then subjected to UV for 30 minutes inside a biosafety cabinet. The cell suspension was made by diluting 300,000 cells in 1 ml of DPBS-0.01% Pluronic solution that was sterile filtered with a 0.2 μm filter (Pall). The cell suspension was loaded to a 1ml syringe (NIPRO) with a UNP-23 precision tip (Unicontrols) connected to a Tygon tubing (0.02-inch inner diameter and 0.06-inch outer diameter). The whole microfluidic system was driven by a two-channel syringe pump (Fusion 200, CHEMTX Inc.). The cells were deformed through the constrictions at a volume flow rate of 100μl/minute for 2 minutes (60,000 cells were deformed for each experiment). The collected cells were spun down using a benchtop microcentrifuge at 6000rpm for 30s and the supernatant was discarded. The resulting cell pellet was resuspended with 200μl complete DMEM and seeded to one well of a 6 -well plate containing 2ml of complete medium for 4 hours. After that, the medium with floating dead cells was discarded and the survived and attached cells were washed with DPBS twice to further wash off the unattached dead cells. The resulting survivor cells were expanded for 4 days before transferring to a T25 cell culture flask (Thermo Fisher). During the transfer, each 6-well plate (cells from 6 different microfluidic channels) was merged into one flask as one independent group. For each experiment in this study, at least 3 such independent groups were generated and tested. The selected cells were cultured to 80% confluency which generally takes 7-10 days after selection before carrying out the subsequent experiments.

### Viability assessment

Calcein-AM/Ethidium Live/Dead kit (Thermo Fisher, Cat#L3224) was used to quantify the survival rate of MCF7 cells after deforming through the microfluidic deformation assays. For all other cell lines, a trypan blue exclusion (Lonza) assay was used for survival rate quantification. Cells coming out from the outlets of the microfluidic devices were collected and spun down before staining with a pre-mixed solution of Calcein-AM and Ethidium per the manufacturer’s instruction. The cells were stained for 5 mins and loaded into a hemocytometer C-chip (DHC-N01, IN-CYTO). The chip was imaged by a Nikon-A1Rsi confocal immediately to detect the calcium AM (488nm laser) and Ethidium (568nm laser) signals in the cells. The obtained images were processed using Imaris 9.5.0 (Bitplane) to determine the number of live/dead cells. For trypan blue exclusion experiments, pelleted cells were resuspended with a pre-mixed staining buffer (DPBS and trypan blue at 1:1 ratio) and manually counted immediately under an optical microscope. To correctly reflect the variation of the survival test, the WT viability tested in the different experiments was merged into one set and shown consistently in the different quantifications here. MTT assay was used for the viability assessment of cells treated with doxorubicin following the standard protocol from the manufacturer.

### Immunofluorescence

For *Suspended cell imaging*. For experiments that imaged the cells right after deformation, the cells were imaged in a suspension state. Briefly, the cells deformed through the device or cells in suspension were fixed with 4% PFA for 10 minutes. The suspension was then quenched with 100mM glycine in DPBS and spun down with a microcentrifuge. The cells were resuspended in DPBS before transferring to a 35mm glass bottom petri-dish (Iwaki) treated with 2% Pluronic for 30 minutes. The cell suspension was left for sedimentation and attached to the bottom of the petri-dish for 30 minutes at room temperature. After cell attaching, they were permeabilized with DPBS containing 0.5% Triton X-100 for 10 minutes, blocked with DPBS containing 0.1% Triton X-100 and 3% BSA for 2 hours. The cells were then incubated with primary antibodies diluted in an antibody dilution buffer (DPBS with 0.1% Tween-20 and 1% BSA) at 4 ºC overnight, washed with PBST 3 times, and incubated with secondary antibodies diluted with the antibody dilution buffer. Antibodies were used with the following concentrations: 1:1000 mouse anti-Lamin A/C (Cell Signaling Technology, Cat#4777S); 1:1000 rabbit anti-Lamin B1 (Cell Signaling Technology, Cat#13435S); 1:500 rabbit anti-γH2Ax (Cell Signaling Technology, Cat#9718); 1:2000 Alexa Fluor 647, 568, 488 goat anti-mouse and anti-rabbit IgG (Abcam, Cat#ab150079, ab150116, ab150113, ab150077). Actin was stained with 1:2000 Alexa Fluor 647-phalloidin (Abcam, Cat#176759), and DNA was stained with Hoechst 33342 (1:1000, 10mg/ml, Invitrogen, Cat#H3570). For *labeling dead cells after deformation*, 0.2μl of propidium iodide (1mg/ml, Invitrogen, Cat#P3566) was added to 200ul of the collected cell suspension and stained for 10 minutes. The cell suspension was then spun down with a microcentrifuge and the staining solution was removed before fixing with 4% PFA. For *adherent cell imaging*, the cells were seeded on an Iwaki glass bottom petri dish for 24 hours before fixing, and all the subsequent steps were the same as suspended cell imaging. For *cells imaged inside the microfluidic channels*, the protocol was similar only that the respective reagents were injected into the channel with a 1ml syringe carefully connected to the channel at a flow rate of 1μl/min.

### Imaging

All *confocal images* presented in this study were captured using a Nikon A1Rsi point scanning confocal microscopy with a Nikon Ti2-E motorized inverted microscopy and Perfect Focus System. Images were taken with an Andor DU897 EMCCD and a 100x objective immersed with oil (CFI Plan ApochromatVC N.A. =1.40). The system was driven by NIS Elements 5.01 (Nikon) software. Time-lapse imaging for cell tracking and proliferation analysis was recorded using an IMQ Biostation with a 37 ºC, 5% CO2 humidified chamber. The imaging interval was 5 mins for 96 hours and images were taken using a 10x objective.

### Image analysis

The intensity profile was analyzed with ImageJ (NIH), and the nuclear lamin intensity with live/dead labeling was analyzed using Imaris 9.5.0 (Bitplane). The nuclear architecture in Fig. 3 was analyzed with a custom-written MATLAB code^[57]^.

### Proliferation Measurement

For video-based single-cell colony formation quantifications, cell numbers were manually counted from the time frame to guarantee the accuracy of the quantification. For cell number tracking at different time points, the cells were harvested through trypsinization and resuspended in 10ml Isoton buffer. The cell numbers were measured using a Multisizer 4e Coulter Counter (Beckman Coulter). In the experiments quantifying the percentage of proliferating cells in Fig. 5, Click-iT Edu assay (Thermo Fisher, Cat#C10337) was used following the kit manual. Briefly, the cells were seeded 50,000/well on a 12-well plate 1 day before the experiment. At the beginning of the experiments, cells were cultured in an Edu-containing medium for 2 hours. Following that, the cells were fixed and labeled per the manufacturer’s instruction. After labeling, the cell nuclei were stained with 10 μg/ml Hoechst 33342 (Cat#H3570) for 10 minutes, washed 3 times, and kept in PBS. The samples were then imaged using a Nikon A1Rsi confocal microscopy and the images were processed using Imaris (Bitplane) to determine the percentage of Edu positive or negative cells.

### Immunoblotting

Cells were trypsinized with 0.05% trypsin-EDTA in DPBS for 10 minutes and quenched by complete-DMEM followed by centrifuging. Then, the cell pellets were lysed in ice-cold RIPA buffer ((50 mM Tris pH 7.3, 0.25 mM EDTA, 150 mM NaCl, 1% Triton X-100, 1% (w/v) Sodium Deoxycholate, supplemented with protease inhibitors) and centrifuged. The protein-containing supernatants were subjected to SDS-PAGE and western blotting. Antibodies used for western blot were: 1:1000 mouse anti-Lamin A/C (Cell Signaling Technology, Cat#4777S); 1:1000 rabbit anti-Lamin B1 (Cell Signaling Technology, Cat#13435S); 1:1000 rabbit anti-GAPDH (Cell Signaling Technology, Cat#5174), 1:2000 HRP goat anti-mouse and anti-rabbit IgG (Abcam, Cat#ab205719, Cat#ab205718).

### Drug treatment

Doxorubicin (Dox) was used in this experiment for testing the DDR and DNA damaging resistance of the MCF7 cancer cells. A same number of ORI/S1/S7 MCF7 cells were cultured in a 96-well plate until 80% confluency. Eight Dox concentrations, ranging from 0 to 3μM, in the DMEM medium were prepared using the serial dilution method with a dilution ratio of 1:3. After treated with Dox for 3 days, cell viability was examined using the MTT assay and a plate reader.

### RNA sequencing and analysis

RNA was extracted from cells harvested by trypsin with RNA extraction kit (Qiagen, RNeasy Mini Kit), and each group was done in duplicates. The extracted RNA samples were shipped with dry ice for sequencing in BGI HK using a DNBseq platform. Reads mapped to rRNA, containing low quality, adaptor-polluted, and high content of unknown base (N) reads were filtered from the raw reads. Clean reads were mapped to the reference genome using HISAT2^[58]^ and 95% of the reads were mapped on average. Reads were aligned to reference using Bowtie2^[59]^ and gene expression levels of each sample were calculated using RSEM^[60]^. Differentially expressed genes between groups were generated using the DESeq2^[61]^ package in R. The derived DEGs were filtered by P<0.05 when compiling a final DEG list for each comparison. The DEG list comparing CTRL and S1 MCF7 cells with log2FoldChange>0.5 and P<0.05 was used for GO analysis^[62]^ and KEGG pathway analysis^[63]^. A gene set enrichment analysis^[64]^ (GSEA) of the MSigDB hallmark gene sets (www.gsea-msigdb.org/gsea/) were performed with the DEGs pre-ranked by fold change.

### Bioinformatic analysis

Survival analysis in a breast cancer patient cohort with microarray data was done using Kaplan Meier plotter^[65]^ (http://kmplot.com/analysis) with a median cutoff and 120-month follow-up. All probes for the same gene were analyzed and genes with log-rank P <0.05 were considered to have prognostic values in the corresponding endpoint. The following datasets were merged for analysis carried out with each endpoint: E-MTAB-365, E-TABM-43, GSE11121, GSE12093, GSE12276, GSE1456, GSE16391, GSE16446, GSE16716, GSE17705, GSE17907, GSE18728, GSE19615, GSE20194, GSE20271, GSE2034, GSE20685, GSE20711, GSE21653, GSE2603, GSE26971, GSE2990, GSE31448, GSE31519, GSE32646, GSE3494, GSE37946, GSE41998, GSE42568, GSE45255, GSE4611, GSE5327, GSE6532, GSE7390, GSE9195.

GSE2603 was compiled using the *GEOquery* package in R for obtaining the differentially expressed genes in parental MDA-MB-231 and lung metastasized MDA-MB-231 breast cancer cells. The expression matrix was compiled and processed with the *limma* package.

TCGA breast cancer patient (BRCA) cohort data were downloaded and compiled into a complete dataset with the expression level of each gene using *TCGA-Assembler*^[66]^. The BRCA patient information and tissue type information are retrieved from the patient barcodes to distinguish metastasis tumors and primary tumors.

Pan-cancer LMNA, LMNB1, and LMNB2 gene expression analyses comparing tumor and normal tissues were performed with the help of Firebrowse (www.firebrowse.org, Board Institute) and the raw data were retrieved from the graph and replotted. All data analyses and plots in this part were carried out using custom codes in R studio.

### Statistics and data presentation

All statistical tests were noted in the legend of each figure. Generally, student’s t-test was used in data with exactly three measurements of each group. Non-parametric test was used in data not following norm distribution with normality checked with Q-Q plot. One-way ANOVA was used in analysis with more than 2 groups. The data range in the figures is represented with mean±s.e.m and jitters showing the data distribution. P values are represented as follows: *** <0.001; ** <0.01; exact values >0.01. The statistical test was performed using the relevant packages with R. All figures in this paper were plotted using R, and mainly with *ggplot2* packages. Survival analysis was performed using the R *surveminer* and survival packages.

## Author contributions

Conceptualization: KJ, CTL

Methodology: KJ, SBL, JX, MS, DSJ, BCL, GVS, CTL

Investigation: KJ, SBL, JX, MS, DSJ, XS, PZ, LL

Visualization: KJ, SBL, JX, XS

Funding acquisition: CTL Supervision: CTL

Writing – original draft: KJ

Writing – review & editing: KJ, SBL, JX, MS, DSJ, XS, PZ, LL, BCL, GVS, CTL

## Data and materials availability

Source data and codes for generating each Figure can be requested from the author upon reasonable request. The RNA-Seq results will be deposited in the Gene Expression Omnibus (GEO, NCBI).

## Supplementary Materials

Figure. S1 to S8

Movies S1 to S5

## Acknowledgments

We thank Dr. Bingxue Yu and Dr. Thuan Beng Saw for substantial discussion of the project, MBI microfabrication core, wet lab core, and microscopy core for helping with the experiments, and Dr. Xing Fei Tan for generating the luciferase-expression cell lines. We also thank Mr. Zac Goh for drawing the illustrations presented in this work.

