## Supplementary material for "Deleterious Mechanical Deformation Selects Mechanoresilient Cancer Cells with Enhanced Proliferation and Chemoresistance": Figure. S

**This PDF file includes:**

Figs. S1 to S8
Captions for Movies S1 to S5

**Other Supplementary Materials for this manuscript include the following:**

Movies S1 to S5

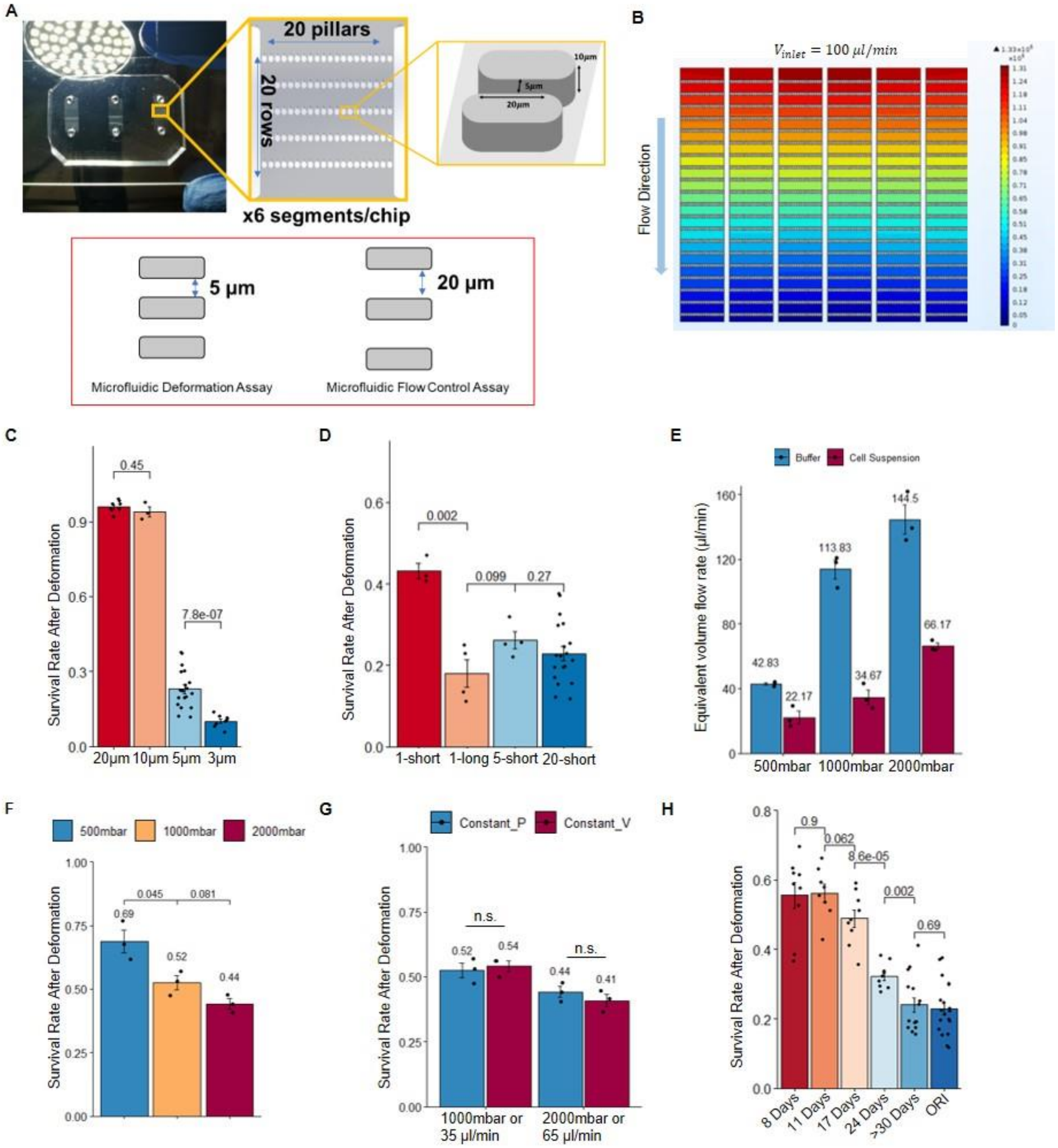


**Fig. S1. Survival rate of cancer cells passing through different configured channels for determining the parameters for the selection experiments. (A)** Top: Schematics of a microfluidic device with multiple rows of small pillars with 5µm gap size which are used for performing mechanical selection on cancer cells. Bottom: illustration of Microfluidic Deformation Assay: pillars have a gap size of 5 µm, and Microfluidic Flow Control Assay: pillars have a gap size of 20 µm. **(B)** COMSOL simulation of pressure distribution inside the microfluidic deformation assay with 100µl/min input flow rate**. (C)** The survival rate of MCF7 cells deformed through 20 rows, 10 µm high channels with varying widths. Barplot showing mean±s.d.; unpaired, two-tailed, t-test with exact p values labeled on the graph, each jitter shows one independent repeat (the same for **(D-H)**). **(D)** The survival rate of MCF7 cells deformed with single or multiple short constrictions, or 1 long constriction. Barplot showing mean±s.d.; unpaired, two-tailed, t-test with exact p values labeled on the graph. **(E)** The equivalent volume flow rate of different inlet pressure at constant pressure infusion mode. Barplot showing mean±s.d. **(F)** The survival rate of MCF7 cells deformed with different input pressure in constant pressure mode. Barplot showing mean±s.d.; unpaired, two-tailed, t-test with exact p values labeled on the graph. **(G)** Comparison of the survival rate of MCF7 cells deformed with constant pressure mode or with the matched volume flow rate in constant flow rate mode. Barplot showing mean±s.d.; unpaired, two-tailed, t-test; n.s.: $p > 0.05$. **(H)** The survival rate of MCF7 cells deformed with microfluidic deformation assay at different time points post mechanical selection. Barplot showing mean±s.d.; unpaired, two-tailed, t-test with exact p values labeled on the graph.

**Figure S2**

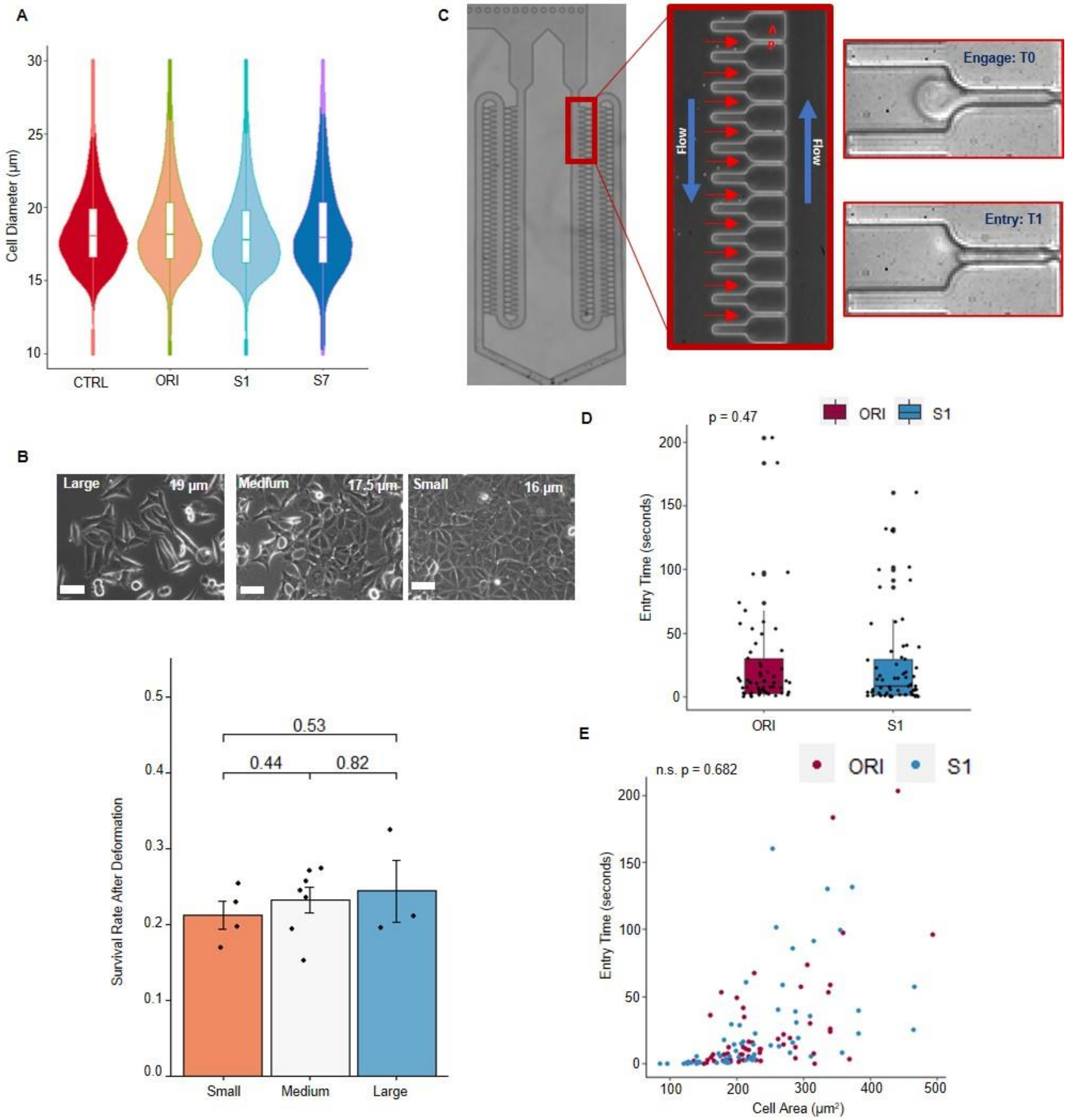


**Fig. S2. Biophysical characterizations of the mechanoresilient MCF7 cells. (A)** Quantification of the cell size in original or selected MCF7 cells; violin plots showing the size distributions, representative distributions from at least 3 independent measurements of each group were shown here. **(B)** The survival rate of MCF7 cells in different size ranges after deforming through the microfluidic deformation assay; Barplot showing mean±s.d; unpaired, two-tailed, t-test with exact p-value labeled on the graph, each jitter shows one independent repeat. **(C)** A microfluidic micropipette assay for assessing the deformability of cells; the time spent from cell engagement to full entry is used to quantify the deformability. **(D)** Comparison of the deformability of ORI and S1 MCF7 cells, n=57 and n=62 in ORI and S1 group respectively; unpaired, two-tailed, t-test with p-value labeled on graph. **(E)** Co-plot of cell area and entry time into the microfluidic micropipette;

generalize linear model with cell area and cell type as covariates; no statistically significant difference of entry time can be seen between the ORI and S1 MCF7 cells.

**Figure S3**

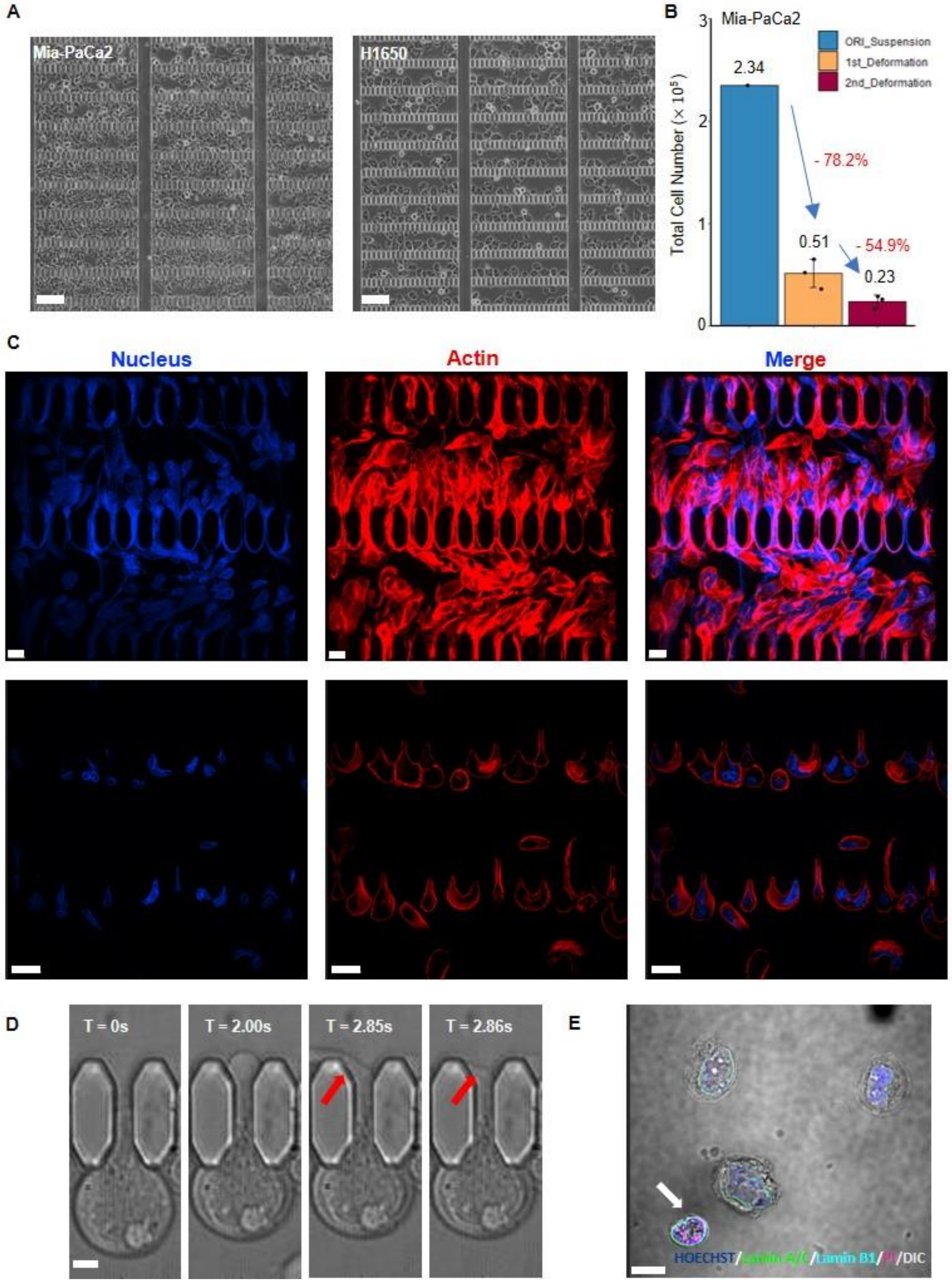


**Fig. S3. Cancer cell behavior during the mechanical selection process. (A)** Representative images of the microfluidic deformation assay after performing mechanical selection experiment on Mia-Paca2 (left) or H1650 (right). Scale bar: 100µm. **(B)** The total cell number in the original cell suspension and cell suspension after each round of deformation (up to two rounds). Barplot showing mean±s.d, n=3. **(C)** Representative images of the microfluidic deformation assay after the mechanical selection experiment. Significant cell ruptures can be seen in the Mia-PaCa2 cells but not in H1650 cells. Scale bar: 20µm (top) and 50µm (bottom). **(D)** The deformation dynamics at the cellular level of an MCF7 cell entering a constriction. A cell body fragmentation is indicated by the white arrows. This phenomenon is only observed in large cells with a diameter>20µm. Scale bar: 5µm. **(E)** A stand-alone nucleus after deformation indicating a complete cell body rupture. Scale bar: 10µm.

**Figure S4**

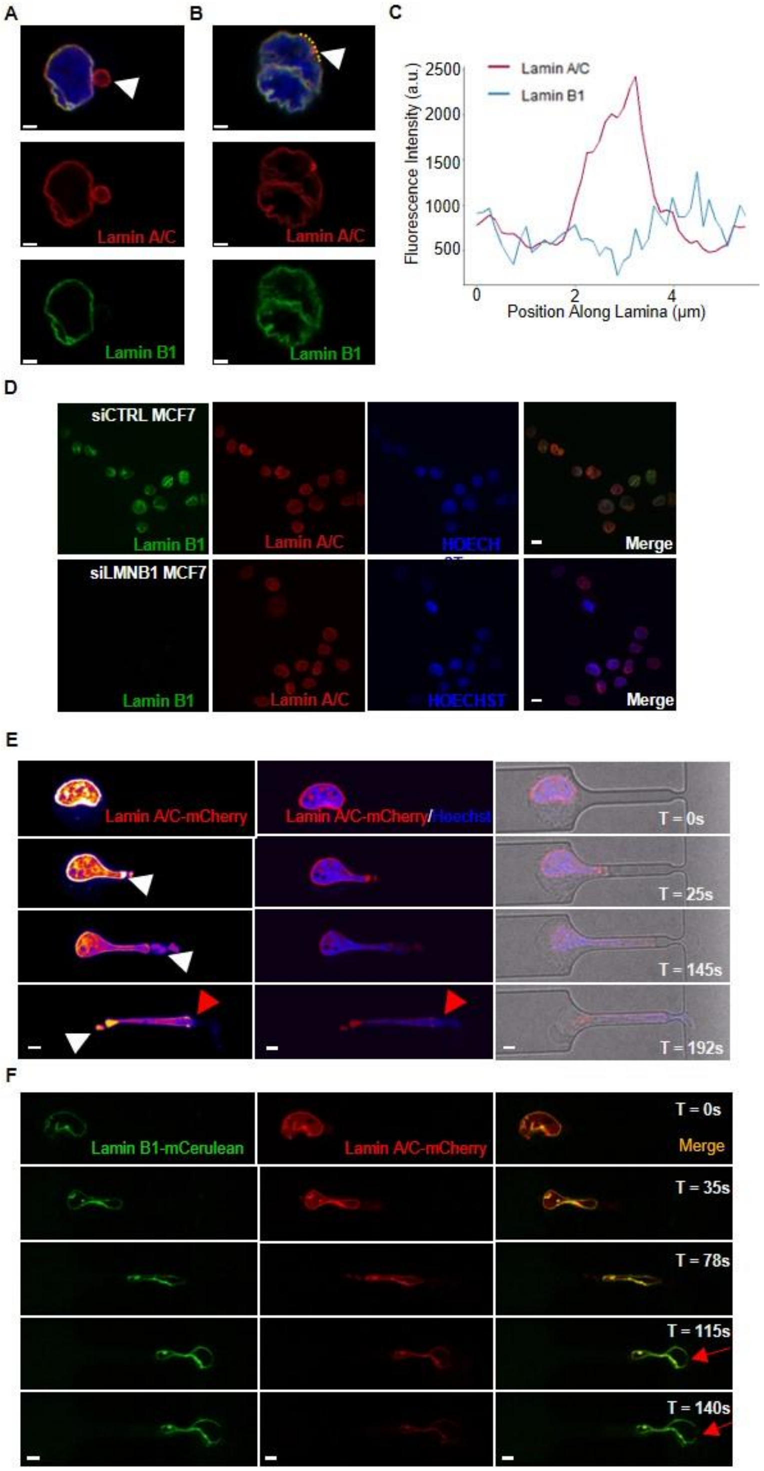


**Fig. S4. Nuclear lamina phenotypes in the survivor cells after deformation or lamin B1 knockdown and the deformation dynamics of the nuclear lamina. (A)** Representative images showing a nuclear blebbing in a survivor MCF7 cell. Scale bar: 4μm. **(B)** Representative images showing a point accumulation of lamin A/C potentially indicating a site of previously damaged nuclear lamina; Scale bar: 4µm. **(C)** The intensity profile of lamin A/C and lamin B1 along the dotted line in **(B)**. **(D)** Representative images showing the nuclear lamina after lamin B1 knockdown. There is no apparent difference in the nuclear shapes of MCF7 cells after lamin B1 gets depleted. Scale bar: 10µm**. (E)** The dynamics of lamin A/C-mCherry when aspirated into a microfluidic micropipette. The left panel shows the intensity of lamin A/C with heat LUT. White arrows pointed to sites where lamin A/C get segregated from the nuclear lamina. Red arrows pointed to the site where lamina ruptures and DNA leaks out. Scale bar: 10 µm. **(F)** A MCF7 nucleus is co-labeled with lamin B1-mCerulean and lamin A/C-mCherry. When deforming into the microfluidic micropipette, lamin A/C gets segregated and weakens while no fluctuation from lamin B1 is observed. Red arrows indicated a site where the lamina ruptured when squeezed through the 2µm constriction at the end of the micropipette. Scale bar: 10 µm.

**Figure S5**

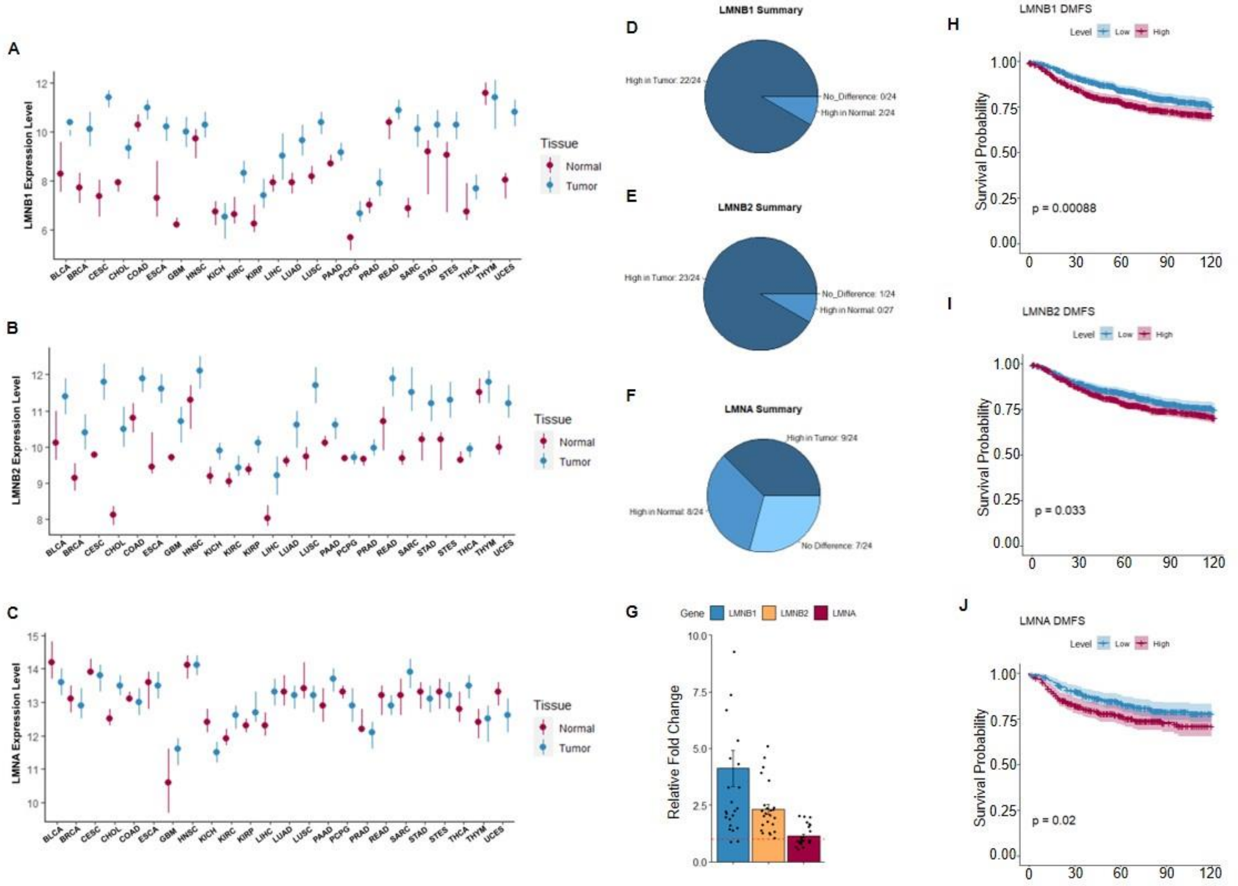


**Fig. S5. Pan-cancer analysis of nuclear lamin expressions and their association to the prognostics of breast cancer. (A-C)** Expression levels of LMNB1, LMNB2, and LMNA in different types of cancer comparing the normal tissues and tumor tissues, point range showing median with 75 and 25 percentiles, data compiled from Firehorse (see Methods). Cancer type full names: BLCA-Bladder Urothelial Carcinoma, BRCA-Breast invasive carcinoma, CESC-Cervical squamous cell carcinoma and endocervical adenocarcinoma, CHOL-Cholangiocarcinoma, COAD-Colon adenocarcinoma, ESCA-Esophageal carcinoma, GBM-Glioblastoma multiforme, HNSC-Head and Neck squamous cell carcinoma, KICH-Kidney chromophobe, KIRC-Kidney renal clear cell carcinoma, KIRP-Kidney renal papillary cell carcinoma, LIHC-Liver hepatocellular carcinoma, LUAD-Lung adenocarcinoma, LUCS-Lung squamous cell carcinoma, PAAD-Pancreatic adenocarcinoma, PCPG-Pheochromocytoma and Paraganglioma, PRAD-Prostate adenocarcinoma, READ-Rectum adenocarcinoma, SARC-Sarcoma, STAD-Stomach adenocarcinoma, STES-Esophagus-Stomach Cancer, THCA-Thyroid carcinoma, THYM-Thymoma, UCEC-Uterine corpus endometrial carcinoma. **(D-F)** Summary of the respective gene levels in tumor or normal tissue in the 24 cancer types analyzed in a-c. **(G)** Summarized average foldchange comparing tumor tissues and normal tissues of the three lamin genes in the 24 cancer types, the red line marked fold change=1. **(H-J)** KM plot of the survival analysis of the three lamin genes using distant metastasis-free survival (DMFS) as the clinical endpoint. Analysis performed in a merged breast cancer database with 664 breast cancer patients (see Methods.)

**Figure S6**

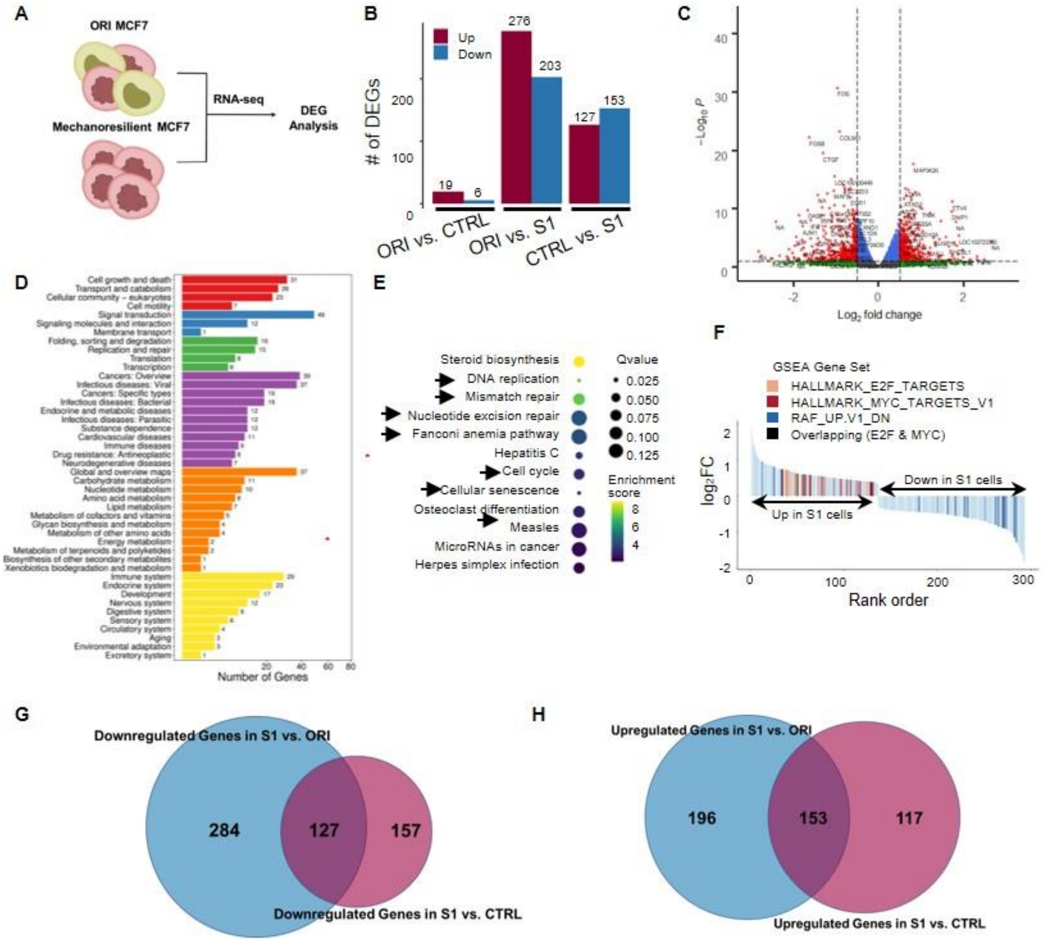


**Fig. S6. Upregulated proliferation and DNA damage repair pathways in mechanoresilient MCF7 cells. (A)** Design of the RNA-Seq experiment. **(B)** Quantification of the DEG numbers paired by different DEG analysis groups, red/blue colors indicated the numbers of up/down-regulated genes. **(C)** Volcano plot of the DEGs comparing the CTRL and S1 cells, log2FoldChange > 0.5, and LogP<-3 are marked as dashed lines in the graph. **(D)** GO enrichment analysis of the CTRL vs. S1 DEG list; red stars highlight the proliferation and DDR-related terms. **(E)** Top enriched KEGG pathways in the CTRL vs. S1 DEGs; arrows marker the pathways related to proliferation and DNA damage repair. **(F)** GSEA of E2F and Myc target genes in the DEG list of CTRL vs. S1 group, each vertical line stands for one gene and the genes are ranked by their expression fold change. **(G)** Common downregulated and **(H)** upregulated genes in S1 vs. ORI and S1 vs. CTRL groups with adjusted p-value < 0.01 as a cutoff.

**Figure S7**

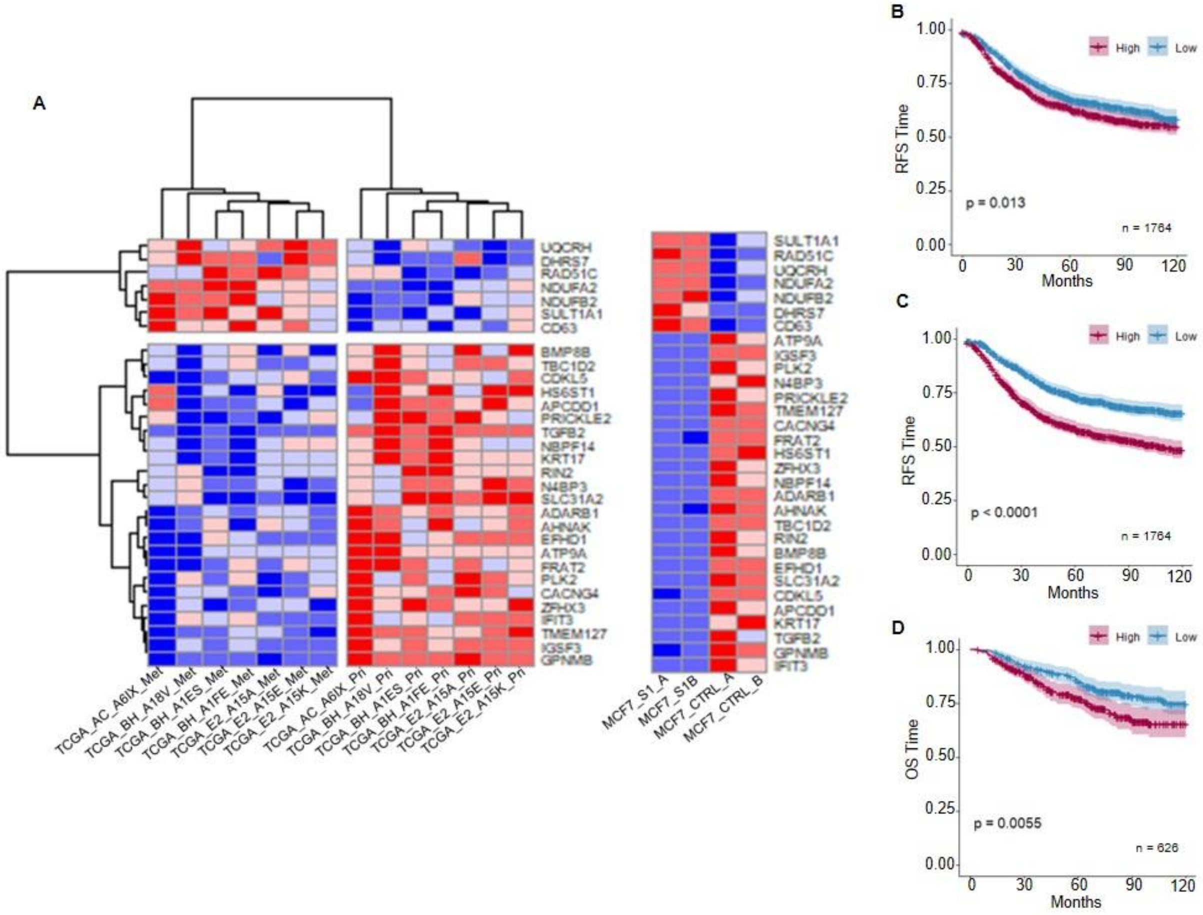


**Fig. S7. Mechanoresilient DEGs are correlated with breast cancer metastasis and prognostics. (A)** Heatmap of 31 genes overlapping between the paired primary and metastasis tumor DEGs in TCGA BRCA cohort and DEGs in S1 MCF7 (adjusted p <0.05 cutoff). **(B)** The KM plot of the 7 metastasis-associated genes in **Fig. 6E-K** using RFS as a clinical endpoint. **(C)** The KM plot of the 31 metastasis-associated genes in **(A)** using RFS and **(D)** OS as clinical endpoints.

**Figure S8**

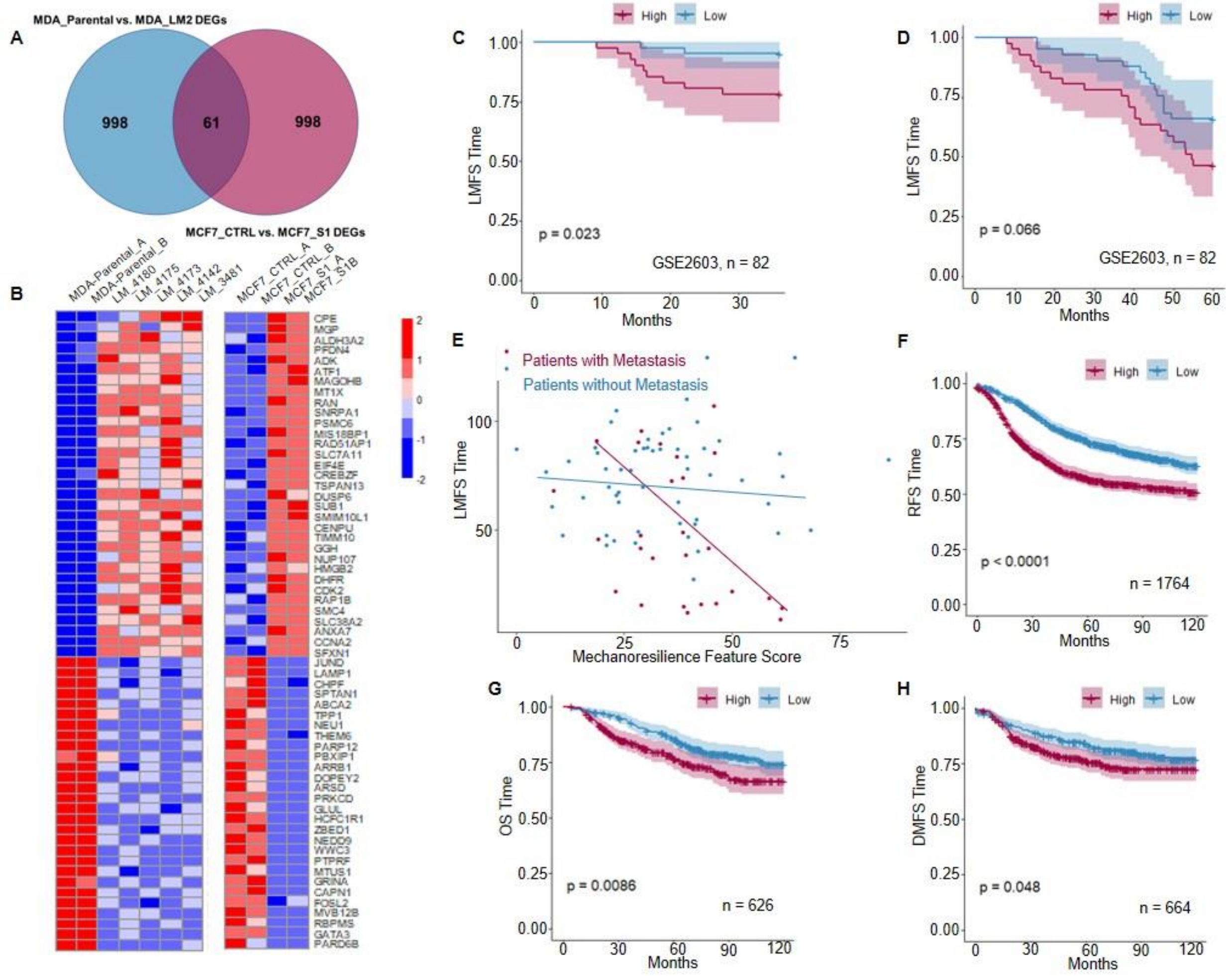


**Fig. S8.** ***In vitro*** **mechanical selection is correlated to the** ***in vivo*** **metastatic selection of breast cancer cells.** **(A)** Overlapping of the DEGs in MDA_LM2 and MCF7_S1. **(B)** Heatmap of the expression of 61 common genes in the two studies in **(A)**. **(C)** 3-year, and **(D)** 5-year lung metastasis-free survival of breast cancer patients in GSE2603 cohort using the weighted expression of the 61 genes as a score. **(E)** The correlation between the score of mechanoresilient signature genes and LMFS with lymph node metastasis status labeling. The score has a much stronger prognostic power in the metastatic subgroup. **(F)** KM plot of RFS, **(G)** OS, and **(H)** DMFS in a merged breast cancer dataset (see Methods).

**Movie Captions**

**Movie S1.** MCF7 breast cancer cells squeezing through the capillary-mimicking constrictions in the microfluidic deformation assay. White arrow indicates the flow direction.

**Movie S2.** Measuring the deformability of a cancer cell using the microfluidic micropipette assay. White arrow shows the flow direction.

**Movie S3.** (**A)** An intact MCF7 cell transiting through the constriction. (**B)** An MCF7 cell transiting with eventual cell rupture. Yellow arrow near the end of the video shows the rupture site.

**Movie S4.** An MCF7 cell transfected with mCherry-LaminA/C (labelling nuclear lamina protein laminA/C) squeezing through a 5μm constriction. The color map shows the relative intensity of mCherry fluorescence.

**Movie S5.** An MCF7 cell labeled with **a,** mCelulean-LaminB1 (labelling nuclear lamina protein lamin B1) and **b,** mCherry-LaminA (labelling nuclear lamina protein lamin A/C) squeezing through a 5μm constriction. The channel opening has width of 2μm.
